# *In-situ* diversification and regional attributes shape asymmetric diversity of *Miliusa* (Annonaceae) in tropical Asia

**DOI:** 10.1101/2025.05.22.655121

**Authors:** Abhishek Gopal, Navendu Page, Amit Kumar, Neha Tiwari, Velusamy Sundaresan, Jahnavi Joshi

## Abstract

**Aim:** We examine biogeography and speciation patterns in *Miliusa* Lesch. ex A. DC. (∼65 species) distributed in tropical Asia to understand the uneven distribution of its extant diversity.

**Location:** Tropical Asia

**Taxon:** *Miliusa* (Annonaceae)

**Methods:** Phylogenetic reconstruction was performed using six plastid markers across 52 species using both ML and BI approaches. Divergence times were estimated using two fossil calibrations and an optimised relaxed clock, and ancestral areas were inferred with BioGeoBEARS. Speciation rates were examined using ClaDS and the DR statistic, and community structure was assessed using phylogenetic diversity metrics.

**Results:** *Miliusa* likely originated in the mid-Miocene, with Indo-Burma and peninsular India (PI) as its ancestral range. Its extant diversity is primarily attributed to *in-situ* speciation, with dispersal or vicariance playing limited but important roles in PI, and Wallacea and Sahul. Lineages in Indo-Burma began accumulating in the mid-Miocene, preceding those in PI (∼10 Myr) and Wallacea and Sahul (∼5 Myr). PI showed signs of lineage saturation and had lower speciation rates compared to Wallacea and Sahul and Indo-Burma, both of which had similar rates. All regions exhibited phylogenetic clustering, but Indo-Burma and PI differed in their sensitivity to phylogenetic depths.

**Main conclusions:** The uneven diversity of *Miliusa* is shaped by time for speciation, age, and dispersal, although their relative influence varies across regions. In Indo-Burma, long-term geo-climatic stability and greater niche availability likely facilitated the persistence of lineages, rapid speciation, and dispersal, making it an evolutionary hotspot for *Miliusa.* In contrast, PI exhibited lower richness and speciation rates despite being old, likely due to the contraction of wet habitats in the Miocene that limited available niches for speciation. Lineages in Wallacea and Sahul show typical island-like radiations, with speciation rates comparable to the larger and more geo-climatically stable Indo-Burma, despite their more recent origin. Overall, our results highlight the role of Miocene-driven climatic vicariance and Pliocene-Pleistocene climatic fluctuations in shaping the diversification dynamics and regional diversity patterns across tropical Asia.

## 1. Introduction

The examination of uneven spatial and temporal distribution of diversity and its drivers is a central theme in biogeography. Broadly, three non-mutually exclusive processes can be invoked to explain spatial heterogeneity in diversity: the time available for speciation, speciation rates, and dispersal rates (Eiserhardt et al., 2017; Igea and Tanentzap, 2019; Ricklefs, 1987; Wiens, 2011). Older and geo-climatically stable regions tend to accumulate more lineages through both *in-situ* speciation and dispersal (Stephens and Wiens, 2003). Alternatively, speciation and dispersal rates can act independently of a region’s age in driving variations in regional diversity patterns (Wiens, 2011). However, the relative influence of these macroevolutionary processes is likely to be modulated by biogeographic history and lineage-specific traits. For example, climatically stable areas act as refugia, allowing for the persistence and accumulation of lineages with similar climatic tolerances through dispersal due to niche conservatism (De Bruyn et al., 2014; Joyce et al., 2020; Skeels et al., 2023). Additionally, these, coupled with larger geographic areas and more heterogeneous habitats, allow for higher speciation due to the availability of a wider array of ecological niches for divergence (Fine, 2015; Schluter and Pennell, 2017).

These macroevolutionary processes have been invoked to explain large-scale diversity patterns, such as the latitudinal diversity gradient (Fine, 2015; Mittelbach et al., 2007) and diversity patterns across biogeographic regions within the tropics (Antonelli et al., 2015; Couvreur et al., 2015; Hagen et al., 2021). However, their examination within a biogeographic region remains limited, particularly in tropical Asia (De Bruyn et al., 2014; Kuhnhäuser et al., 2025). Lineage level examination of these patterns and their drivers has been particularly insightful (Koenen et al., 2015; Li et al., 2024), however, it also remains limited in tropical Asia.

Tropical Asia is one of the most geologically dynamic and speciose regions globally (Lohman et al., 2011), encompassing up to six biodiversity hotspots (Myers et al., 2000). The region’s complex interplay of geological and climatic processes has shaped its biotic assembly (Kooyman et al., 2019; Lohman et al., 2011), resulting in high levels of endemism and species richness that rival those of the Neotropics (Raven et al., 2020). Tectonic collisions during the Cenozoic, involving the Indian and Australian plates with Asia, played a pivotal role in shaping the region’s geology, climate, and biotic assembly, setting the stage for the origin and evolution of rainforests (Kooyman et al., 2019; Lohman et al., 2011; Morley et al., 2018). The collision of the Indian plate in the Eocene triggered the upliftment of the Himalayas and the onset of the monsoon, driving the contraction of wet forests during the Miocene (Morley et al., 2018). Similarly, the collision of the Sahul shelf in the late Oligocene–early Miocene restructured the Sunda–Sahul region, resulting in the emergence of islands and mountains in Malesia (Hall 2009, 2011), and also initiated the East Asian monsoon, creating wetter conditions that supported the establishment of modern Malesian flora (Morley et al., 2018). These geological events enabled the extensive dispersal of rainforest lineages across these regions (Crayn et al., 2014; Joyce et al., 2020; Zhang et al., 2022), especially from the mid-Miocene onwards, facilitated by the proximity of Sunda and Sahul, the emergence of islands in Wallacea and the upliftment of parts of New Guinea (Toussaint et al., 2014). These geological processes, coupled with the Pliocene-Pleistocene climatic oscillations, resulted in recurrent overland connections and disconnections as well as repeated contractions and expansions of wet rainforests (Hall 2009, 2011; Lohman et al., 2011), further influencing the rate of dispersal and diversification of rainforest lineages. Collectively, these geo-climatic dynamics have played a pivotal role in shaping the region’s tremendous diversity and endemism (Kooyman et al., 2019)

However, the interplay of these processes and their impact varied across tropical Asia, driving the disparity in diversity among its biodiversity hotspots. Indo-Burma and Borneo are considered major evolutionary hotspots owing to their relative geological and climatic stability, especially during the Pliocene-Pleistocene, which likely allowed them to serve both as refugia for rainforest lineages and as sources of dispersal to other regions (De Bruyn et al., 2014; Joyce et al., 2020; Kuhnhäuser et al., 2025; Lohman et al., 2011). In addition to these dispersal dynamics, the upliftment and orogeny on islands in Wallacea and Papua New Guinea allowed for extensive *in-situ* speciation of lineages in these regions during the Pliocene-Pleistocene (Joyce et al., 2020; Lohman et al., 2011).

In the context of these biogeographic dynamics, the role of peninsular India (PI) remains unexplored, largely due to a lack of molecular data from the region, specifically for woody plants. While its significance as a “biotic ferry” and its role in spurring the diversification in continental Asia during the Eocene are well documented (Kooyman et al., 2019; Morley et al., 2018), its role in shaping regional diversity from the Miocene onwards remains poorly understood. However, existing evidence suggests that much of the extant flora in PI originated through dispersal from southeast Asia, with most of the relictual flora likely having faced extinction (Morley et al., 2018; Mishra et al., 2024; Parmar et al., 2023; Prasad et al., 2009).

Here, we examine the relative role of these macroevolutionary processes in shaping the uneven diversity in tropical Asia using *Miliusa* Lesch. ex A. DC. belonging to the tribe Miliuseae of the family Annonaceae (Chatrou et al., 2012). Around 65 species of *Miliusa* are currently described (Kew Plants of the World; https://powo.science.kew.org/), and the genus is distributed across tropical Asia, ranging from the Indian subcontinent, the Himalayas, southern China, Indo-Burma, Sundaland, Papua New Guinea, and Australia. The species richness patterns of *Miliusa* show that Indo-Burma is the centre of its diversity, with the highest number of species and endemics being reported from the region. *Miliusa* are mostly shrubs or small trees, primarily occurring in wet habitats and are dispersed by birds and mammals. Previous systematic studies, based on both morphological and molecular data, have identified four clades within *Miliusa* (Chaowasku et al., 2013; Damthongdee et al., 2023). However, only four of the thirteen species from PI (two endemic and two widespread) have been included in these studies, and the divergence time for the genus has not been estimated.

Here, using a densely sampled dataset that covers ∼80% of the *Miliusa* species, including the endemic PI species, and spanning the genus’s entire geographical range, we reconstruct a time-calibrated phylogeny to examine its evolutionary origin, historical biogeography, and speciation patterns. We examine the relative role of time available for speciation, biogeographic processes (dispersal and vicariance), and variation in speciation rates in shaping the current uneven diversity within *Miliusa* across tropical Asia. Furthermore, to assess the signature of these biogeographic and diversification processes on regional assemblages across tropical Asia we examine their phylogenetic structure. A clustered assemblage is indicative of *in-situ* speciation driven assembly, while an overdispersed assemblage is likely due to the effect of dispersal. Lastly, as a case study, we explore the potential role of climatic niches in driving the speciation and distribution patterns using the well-sampled endemic PI clade.

## 2. Materials and Methods

### 2.1. Taxon and DNA sampling

Taxon sampling included 52 species (80% of the 65 accepted species) and 66 accessions of *Miliusa*. From PI, we have near complete sampling with 13 species represented, of which 11 are endemic, and two are widespread. Although 16 endemic species are reported from the region (KEW POWO), five of these are likely to be synonyms (Table S1). Primary data was generated for 31 accessions representing 20 species (12 accepted and 8 unique morphotypes; Table S2), spanning the Western Ghats, PI, the Eastern Ghats, and the Andaman Islands. The primary data was complemented with accessions from the National Centre for Biotechnology Information (NCBI, http://www.ncbi. nlm.nih.gov; Table S2).

Five plastid markers—*mat*K, *ndh*F (*ndh*F1 and *ndh*F2), *ycf*1 exon, *psb*A-*trn*H spacer, and *trn*LF (*trn*L intron and *trn*L-F spacer)—(∼4300 bp) following Chaowasku et al., (2012), were used for the phylogenetic reconstruction and divergence time estimation. For both phylogenetic reconstruction and divergence time estimations, the taxon sampling was extended to all the genera within Annonacea having at least one species representative per genus, and with seven species from the related Magnoliid families as outgroups (Degeneriaceae, Eupomatiaceae, Himantandraceae, Magnoliaceae, and Myristicaceae) following Chen et al., (2019) and Thomas et al., (2017). The final data matrix consisted of 175 accessions, comprising 65 accessions of *Miliusa*, 103 accessions representing other genera from Annonaceae, and 7 outgroups from Magnoliales. DNeasy Plant Mini Kit (Qiagen, Hilden, Germany) and a modified cetyltrimethylammonium bromide (CTAB) protocol (Doyle and Doyle, 1987) were used for DNA extraction from dried leaf samples. PCR amplifications were done following Chaowasku et al., (2012), and the PCR products were sequenced at an in-house Sanger sequencing facility at CSIR-Centre for Cellular and Molecular Biology (CCMB), Hyderabad (Table S3). Sequences were manually cleaned and aligned using MUSCLE (Edgar, 2004) and concatenated in MEGA (Kumar et al, 2016).

### 2.2. Phylogenetic analysis

Phylogenetic reconstructions were done using the maximum likelihood (ML) and Bayesian inference (BI) approaches. The ML tree was run using the command line version of IQTree v.2.3.4 (Minh et al., 2020) using standard bootstrap with 1000 replicates and the built-in model finder (Chernomor et al., 2016; Kalyaanamoorthy et al., 2017) for the concatenated dataset. For BI analyses, the best partition scheme was determined using PartitionFinder2 (Lanfear et al., 2017). The best scheme identified four partitions, with GTR being the best model for *mat*K and *trn*LF, *ndh*F1, *psb*A-*trn*H, and GTR+I being the best model for *ndh*F2 and *ycf*1. The BI was run in MrBayes v.3.2.7 (Ronquist et al., 2012), with four independent Monte Carlo Markov Chain (MCMC) analyses run for 150 million generations and sampled every 5000th generation until the standard deviation of split frequencies was less than 0.05. Trace files were examined using Tracer v.1.7.2 (Rambaut et al., 2018), and the effective sample size (ESS) values for each parameter were checked to see if it was greater than 200. The first 25% of the trees were discarded as burn-in, and strictly bifurcating trees were summarised.

### 2.3. Divergence time estimation

Calibrations were done using two fossils, *Endressinia brasiliana (*c. 112 million years ago, Myr), a fossilised flowering branch, considered to be part of the Magnoliinea crown group and *Futabanthus asamigawaensis* (c. 89 Myr), a fossilised flower, assigned to the Annonaceae crown group, following Pirie and Doyle (2012) and Thomas et al., (2017). The Magnoliinea crown node was calibrated using a log-normal prior distribution, with the age of the *E. brasiliana* as the offset (mean: 10.6, log(SD): 0.6; median: 121.6 Myr, 95% probability interval: 136.4–112.6 Ma; offset: 112.6). Similarly the Annonaceae crown node was constrained using a log-normal prior distribution with the age of *F. asamigawaensis* as the offset (mean 10.7; log(SD):0.6; median: c. 98 Myr, 95% probability interval: 113–89 Ma; offset: 89).

Bayesian divergence time estimation was done in BEAST v.2.7.6 (Bouckaert et al., 2014) using four partitions defined by the partition finder results (see section 2.2). An optimised relaxed clock model (Douglas et al., 2020) with a Yule tree prior was applied. Four independent MCMCs were run for 500 million generations, with every 10,000th generation being sampled. Tracer v.1.7.2 was used to check the MCMC files for convergence, adequate ESS for each parameter, and the burn-in proportion. The tree files of the runs were combined using LogCombiner v.2.7.6 (Bouckaert et al., 2014), resampled at every 20,000 states and discarding 25% as burn-in. TreeAnnotator v.2.7.6 (Bouckaert et al., 2014) was used to summarise the trees post burn-in as maximum clade credibility (MCC) trees with median node heights and discarding 10% as burn-in. Monophy constraints were placed to specify a sister relationship between Myristicaceae and Magnoliineae to avoid erroneous root recovery (Pirie and Doyle, 2012). Furthermore, monophyly constraints were included in the ingroup to constrain taxonomic relationships according to the well-supported nodes from the BI and each species was represented by a single accession.

### 2.4. Ancestral area reconstruction

Three regions were delimited within tropical Asia based on the distribution of extant species and endemicity: PI (A), Indo-Burma (B), which includes the Himalayas, North-east India, continental Asia, and Sundaland, and Wallacea and Sahul (C), including Wallacea, Papua New Guinea and Australia. Ancestral ranges were inferred using BioGeoBEARS (Matzke, 2013, 2014) in R v.4.2.3 (R Core Team, 2023). It provides a likelihood framework to examine range shifts along all branches in the phylogeny, either at the time of branching: cladogenetic events (such as vicariance, founder-event speciation or jump dispersal, and sympatry) or along the branch: anagenetic events (such as dispersal or range expansion and extinction). Six standard models, each with a ‘*j*’ parameter (Matzke, 2014), were examined in an unconstrained framework. Dispersal, extinction, and cladogenesis (DEC; Ree and Smith, 2008), dispersal–vicariance (DIVA; Ronquist, 1997), and a likelihood version of Bayesian biogeographical inference (BAYAREALIKE; Landis et al., 2013). The maximum number of ancestral areas per node was given as two, based on extant species distribution. The best model was selected based on the log-likelihood values and corrected Akaike information criterion (AICc) values (Burnham and Anderson, 2004). All models were tested with the MCC tree estimated from BEAST, and the best-fitting model was identified. To account for uncertainty in both divergence times and tree topology, ancestral ranges (based on the best-fitting model from the MCC) were estimated on 100 trees by resampling from the posterior distribution of the BEAST run following Magalhaes et al., (2021). Additionally, to account for uncertainty in the stochastic mapping, twenty biogeographic stochastic mappings (BSMs; Dupin et al., 2017) were run for each of the hundred trees. BSMs allow quantification of the total number of biogeographic events, their phylogenetic position and directionality. The events were summarised across all 2,000 biogeographic histories to obtain a conservative measure of the cladogenetic and anagenetic speciation events and their transitions.

Further, we also examined how the relative rates of *in-situ* speciation and dispersal (immigration and emigration) differed between the three regions over time. For each BSM replicate, the number of events in each 1 Myr time bin was divided by the number of lineages at the start of the time bin. These rates were then summarised (across 2,000 biogeographic histories) to obtain the median rates and the lower 2.5% and upper 97.5% quantiles (see Magalhaes et al., 2024 for a similar approach).

Lastly, we examined how lineages have accumulated in each of the three regions in 1 Myr time bins using “lineage through time and space plots” (LTST; Skeels et al., 2023), extending it to incorporate topological uncertainty, in addition to uncertainty in the ancestral range estimation following Magalhaes et al., (2021). Lineages in an area can increase via *in-situ* speciation and/or dispersal and decrease via extinctions. Here, based on the 2,000 biogeographic histories, we examined the tempo of lineage accumulation in each of the regions in 1 Myr time bins. In the case of dispersal, we examined the rate of immigration and emigration for lineages in each of the three regions. Range-expansion events and founder-event speciation events were merged to get the overall rates of lineages coming into and dispersing out of a region.

### 2.5. Geographic variation in speciation rates across biogeographic subregions

To examine how speciation rates vary across biogeographic regions, we used two tip-based approaches: a non-parametric approach, the DR statistic (Jetz et al., 2012) and a parametric approach, the cladogenetic diversification rate shift analysis (ClaDS) implemented in the Julia package PANDA v.1.6.1 (Maliet et al., 2019; Maliet and Morlon 2022). Tip rates have been extensively used to compare the patterns of geographic speciation. They represent the best estimate of a lineage’s current speciation rate based on its recent evolutionary history and can be interpreted as an indicator of how soon further speciation might occur (Title and Rabosky, 2018). The DR statistic measures species-specific speciation rates by incorporating the number of splits and branch lengths from the tip to the root, giving a higher weightage to the terminal branches. ClaDS can detect frequent small shifts in speciation rates, which are inherited at the event of speciation, with the daughter lineages being assigned a rate based on a distribution parameterised on the parental rate (Maliet and Morlon, 2022). For ClaDS, MCMC chains were run discarding the first 25% as burn-in and model convergence was assessed by ensuring Gelman statistics were below 1.05 following Maliet and Morlon (2022). A sampling fraction of 0.8 was specified to account for missing species in the phylogeny, given that 80% of the species were included in the phylogeny. Speciation rates were calculated for the entire tree and then extracted for each biogeographic region. Both DR and ClaDS estimates were run on the 100 trees sampled from the posterior distribution (see section 2.4).

### 2.6. Phylogenetic assemblage structure across biogeographic subregions

To examine the relative influence of dispersal versus *in-situ* speciation on the phylogenetic assemblage of *Miliusa* in each region, two community phylogenetic indices were used, mean pairwise distance (MPD) and phylogenetic diversity (PD; Faith, 1992; Webb et al., 2002) along with their respective standardised measures (MPDses and PDses). The standardised measures were calculated using the null model “taxa.labels” in the R package “picante” (Kembel et al., 2010) using 1000 randomisations. Values greater than 1.96 indicate overdispersion, where species are more likely to be distantly related to each other in the assemblage, and values less than -1.96 indicate clustering, where species are more likely to be closely related to each other in the assemblage, while values between -1.96 and 1.96 are considered random. Here, overdispersion is likely to be indicative of a dispersal-driven assemblage structure, while clustering reflect a greater role of *in-situ* divergence in shaping the assemblage structure, with MPDses being sensitive to deeper branching patterns in the phylogeny and PDses to more terminal branching patterns (Mazel et al., 2016). These estimates were calculated for the same 100 trees sampled from the posterior as detailed above (see section 2.4) to account for topological uncertainty.

### 2.7. Climatic niche space occupied by the endemic PI species

To understand the processes driving speciation within a region, we used the PI community for which detailed geographic distribution data are available. We examined the climatic niche space occupied by *Miliusa* in PI (*N* = 13 species; 11 endemic and 2 widespread species) by curating occurrence locations (mean (min-max); 5.69 (1‒14)) from primary collections and through published sources. Three environmental predictors, elevation (m), mean annual precipitation (mm), and precipitation seasonality, were extracted from the WorldClim database (Fick and Hijmans, 2017) to characterise the climatic niche space occupied by *Miliusa*. These predictors have shown to be important determinants of woody plant distribution in the Western Ghats (Page and Shanker, 2020).

## 3. Results

### 3.1. Phylogenetic analysis

We found strong support for the monophyly of *Miliusa* (node: 53; Bayesian posterior probability/ML bootstrap, PP/ML: 1/100%; Fig. S1) and recovered similar clades as previously reported by Chaowasku et al., 2013 with strong support, except for Clade A (Fig. S1). *M. tomentosa, M. velutina*, and *M. sclerocarpa* formed Clade A, where *M. velutina* and *M. sclerocarpa* were sisters with strong support (PP/ML: 1/95%), although the sister relationship of *M. tomentosa* with *M. velutina* and *M. sclerocarpa* had low support (node: 102; PP <0.5; Fig. S1). Lineages from Indo-Burma were part of multiple clades, Clade B2 (node: 80), Clade C (node: 55), and within Clade D (node: 66), respectively (PP/ML: 1,1, and 1/75%, 98%, and 98%; Fig. S1). Clade B (node: 78; PP/ML: 1/92%) comprised of two monophyletic groups: Clade B2 (node: 80; PP/ML: 1/75%) comprising of primarily an Indo-Burman clade and Clade B1 (node: 92; PP/ML: 0.74/<50%) comprising of primarily PI endemics (node: 93; PP/ML: 0.74/<50%), all of which were sister to *M. macrocarpa* from Bhutan. The monophyly of Clade B1 and B2 was not well supported (node: 79; PP/ML 0.79/<59%), with *M. nilagirica,* an endemic species from PI, being recovered as sister to Clade B1 and B2 (PP/ML: 1/92). Clade C (node: 55, PP/ML: 0.99/98%) and Clade D (node: 65, PP/ML: 0.99/99%) were retrieved as sisters with strong support (node: 54, PP/ML: 1/92%). While Clade C had primarily species from Indo-Burma, Clade D comprised two subclades, one with species from Indo-Burma (node: 66, PP/ML: 1/98) and the other with species from Wallacea and Sahul, respectively (node: 73; PP/ML: 1/93%).

### 3.2. Divergence time estimates

Divergence estimation suggests that *Miliusa* originated in the Middle to Late Miocene ∼12 Myr (95% HPD: 15.05–9.03 Myr; node: 53; Table 1; Fig. 1). The median age of Clade A was estimated as 8.71 Myr (95% HPD: 11.82‒5.82 Myr; node: 102). The median ages of Clade B, Clade B1 and Clade B2 were 8.51 Myr, 7.02 Myr and 6.65 Myr, respectively (95% HPD: 11.02‒6.39 Myr, 9.18‒5.12 Myr, and 8.75‒4.81 Myr, respectively; node: 78, 92, and 80). Lastly, the median ages of Clade C and Clade D were 7.07 Myr and 8.05 Myr, respectively (95% HPD: 9.91‒4.75 Myr and 11.01‒5.51 Myr, respectively; node: 55 and 65). A summary of the divergence time estimates (median age and 95% HPD) for key nodes is presented in Table 1, and the MCC tree for *Miliusa* is shown in Fig. 1.

**Fig 1.**
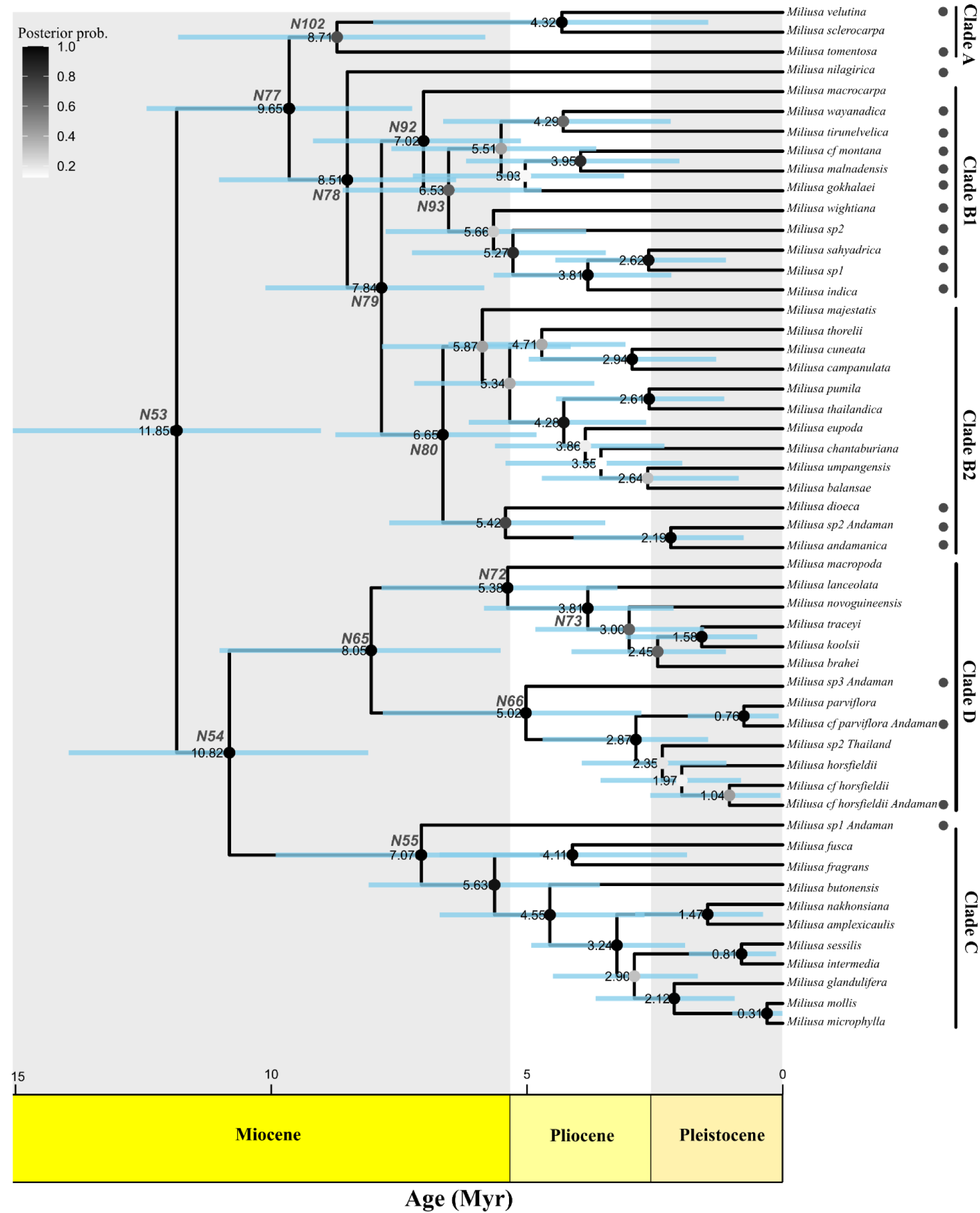
Maximum clade credibility tree of *Miliusa* with the bars representing 95% highest posterior density (HPD) intervals and the nodes representing median ages. The colour of the circle at each node indicates the posterior probability values. Select key nodes are labelled, see Table 1. Clades have been labelled following Chaowasku et al., (2013). Filled circles next to the species names indicate the species for which molecular data was generated in this study.

**Table 1:**
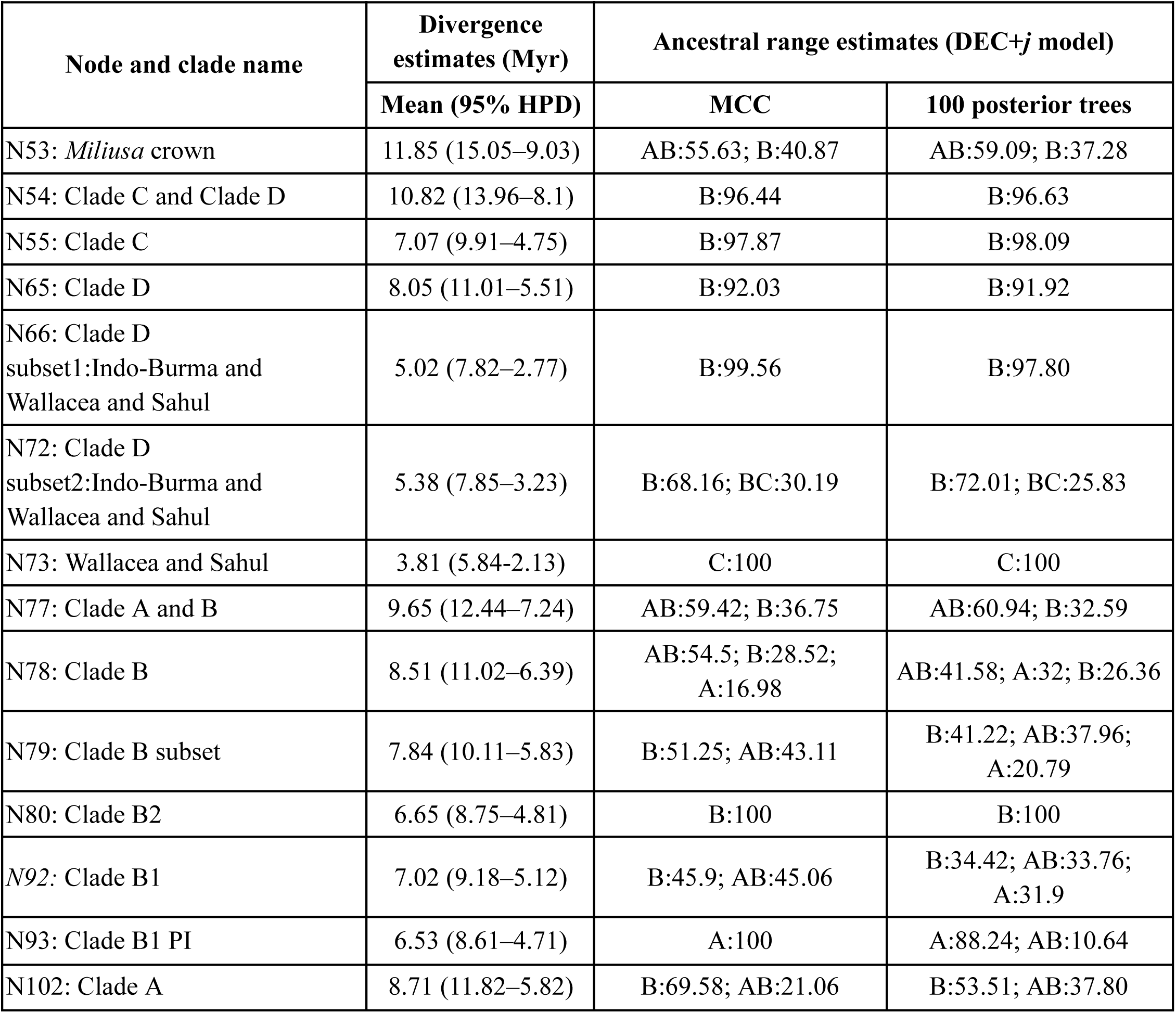
Divergence time and ancestral range estimates of key nodes of *Miliusa* based on the MCC tree and summarised from the 100 trees sampled from the posterior. For the trees sampled from the posterior, only ancestral states with probability values greater than 10% are shown.

| Node and clade name | Divergence estimates (Myr) | Ancestral range estimates (DEC+j model) |  |
| --- | --- | --- | --- |
|  | Mean (95% HPD) | MCC | 100 posterior trees |
| N53: *Miliusa* crown | 11.85 (15.05–9.03) | AB:55.63; B:40.87 | AB:59.09; B:37.28 |
| N54: Clade C and Clade D | 10.82 (13.96–8.1) | B:96.44 | B:96.63 |
| N55: Clade C | 7.07 (9.91–4.75) | B:97.87 | B:98.09 |
| N65: Clade D | 8.05 (11.01–5.51) | B:92.03 | B:91.92 |
| N66: Clade D subset1:Indo-Burma and Wallacea and Sahul | 5.02 (7.82–2.77) | B:99.56 | B:97.80 |
| N72: Clade D subset2:Indo-Burma and Wallacea and Sahul | 5.38 (7.85–3.23) | B:68.16; BC:30.19 | B:72.01; BC:25.83 |
| N73: Wallacea and Sahul | 3.81 (5.84–2.13) | C:100 | C:100 |
| N77: Clade A and B | 9.65 (12.44–7.24) | AB:59.42; B:36.75 | AB:60.94; B:32.59 |
| N78: Clade B | 8.51 (11.02–6.39) | AB:54.5; B:28.52; A:16.98 | AB:41.58; A:32; B:26.36 |
| N79: Clade B subset | 7.84 (10.11–5.83) | B:51.25; AB:43.11 | B:41.22; AB:37.96; A:20.79 |
| N80: Clade B2 | 6.65 (8.75–4.81) | B:100 | B:100 |
| N92: Clade B1 | 7.02 (9.18–5.12) | B:45.9; AB:45.06 | B:34.42; AB:33.76; A:31.9 |
| N93: Clade B1 PI | 6.53 (8.61–4.71) | A:100 | A:88.24; AB:10.64 |
| N102: Clade A | 8.71 (11.82–5.82) | B:69.58; AB:21.06 | B:53.51; AB:37.80 |

### 3.3. Ancestral range estimation

Among the models tested, DEC+*j* had the lowest AICc score, however, the other models (DEC, DIVA and DIVA-like+*j)* were within 2 ΔAICc values (Table S4). While the estimated ancestral states were qualitatively congruent across these models, we proceeded with the DEC models as DIVA does not allow for subset sympatric speciation (Ronquist and Sanmartin, 2011), which, given the biogeographic context, could be plausible for *Miliusa*. Within the DEC models, we selected the DEC+*j* model (Matzke, 2014), wherein new lineages could be established in a new area without the intermediate widespread ancestor. This is a plausible scenario given the multiple island-mainland configurations in tropical Asia and *Miliusa* being dispersed by birds and bats. However, both DEC and DEC+*j* show congruent results on the MCC tree (Table S5), except in three cases where DEC+*j* inferred founder-event speciation while DEC inferred allopatry or subset-sympatry. For these cases, we report and discuss the results of both the models. Additionally, there was a congruence in estimates between the ancestral estimations run on the MCC tree and the average ancestral estimates from the 100 trees sampled from the posterior based on DEC+*j* (Table 1).

According to both the DEC and DEC+*j* models, the ancestral range of the crown node of *Miliusa* (node: 53) was likely widespread, involving areas PI and Indo-Burma (AB) or Indo-Burma (B) alone (Table 1; Fig. 2; Table S5). The early evolutionary history of *Miliusa* is characterised by subset-sympatry primarily from a widespread ancestral range across PI and Indo-Burma (AB) followed by *in-situ* speciation in Indo-Burma (B). Indo-Burma is primarily characterised by *in-situ* speciation (node: 55, 66, and 80). The Wallacea and Sahul region were governed primarily by a founder-event speciation event from Indo-Burma (B → C; node: 72 → 73) or vicariance according to the DEC model (BC → B, C; node 72 → 73), followed by subsequent *in-situ* speciation (Fig. 2; Table S5). Similarly, the PI clade (node: 93) likely originated via either founder-event speciation from Indo-Burma (B → A, B node: 92 → 93, *M. macrocarpa*) or through vicariance (climatic) from Indo-Burma and PI (AB → A, B), followed by *in-situ* speciation. The most recent common ancestor (MRCA) of the Clade B subset (node: 79) is likely to have occupied either AB or B and as such, the descendent nodes either underwent subset sympatry (AB → B, AB; node: 79 → 80, 92) or underwent *in-situ* speciation or narrow sympatry (B → B, B; node: 79 → 80, 92). Lastly, there were two range expansion events from Indo-Burma to PI (node: 102; *M. velutina* and *M. tomentosa*), and from Indo-Burma to Wallacea and Sahul (*M. horsfieldii* and *M. cf. horsfieldii*) and one founder-event speciation from Indo-Burma to Wallacea (*M. butonensis*). While the position of *M. tomentosa* was uncertain, running the DEC+*j* over 100 trees inferred Indo-Burma (B) as its likely ancestral area.

**Fig 2.**
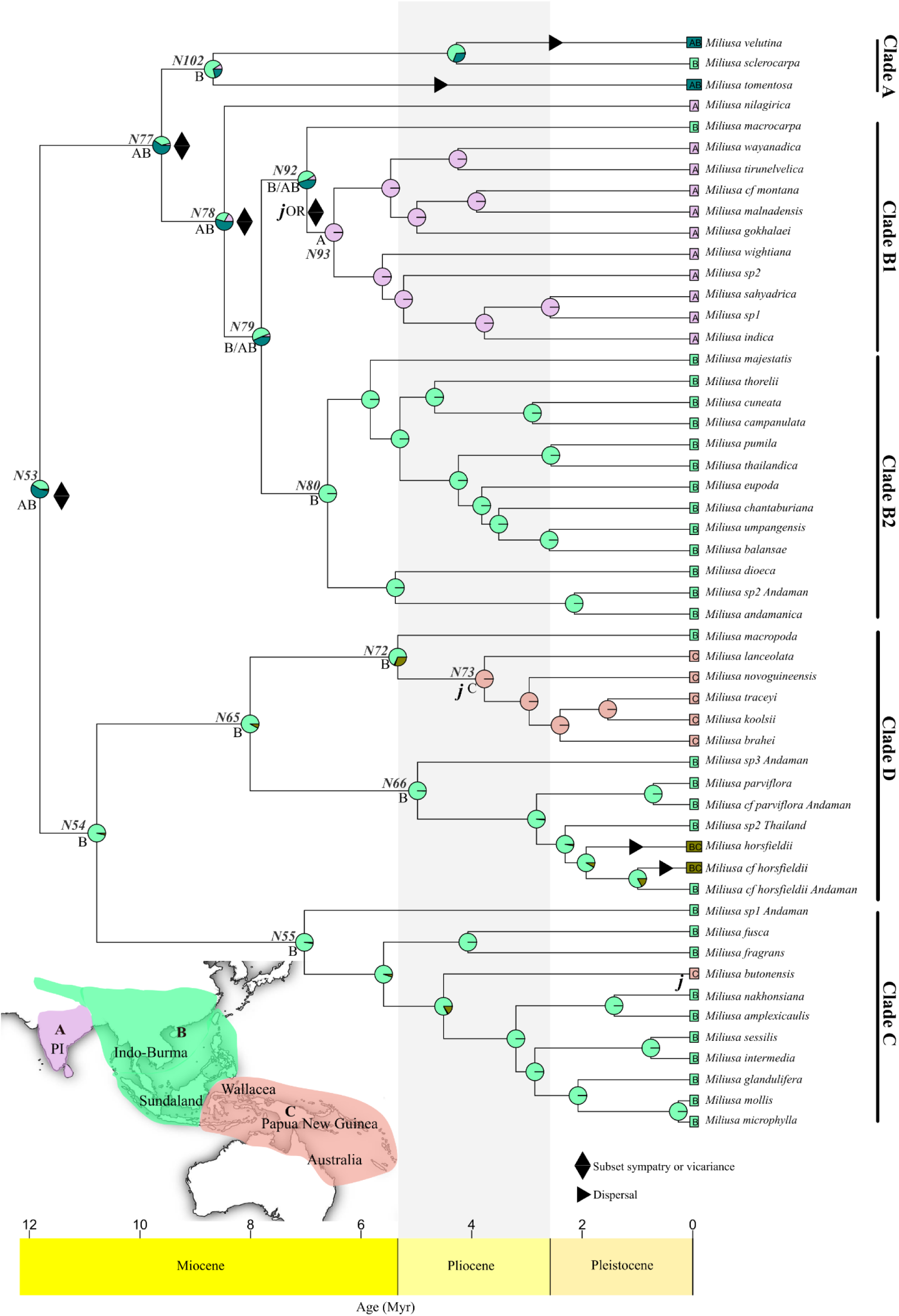
Likelihood-based ancestral range estimation for *Miliusa.* Pie charts at each node indicate the probable ancestral area derived from the DEC+*j* model on the MCC tree. Select nodes are highlighted and correspond with Fig. 1 and Table 1. The text on each node indicates the ancestral state with the highest probability. Areas used in the reconstruction are A: peninsular India (PI), B: Indo-Burma (including the Himalayas, subtropical China, mainland Asia, and Sundaland), and C: Wallacea and Sahul shelf (including Papua New Guinea and Australia). Anagenetic dispersal events, cladogenetic events of subset sympatry, and founder-event speciation or jump dispersal (*j*) are indicated.

After accounting for topological and stochastic mapping uncertainty, our results indicate that overall, within *Miliusa*, 89% of range inheritance events occurred via *in-situ* speciation, followed by subset-sympatry ∼6%, range-expansion ∼5%, founder-event speciation ∼3%, and allopatric speciation ∼2% (Fig S2). These trends were mirrored within Indo-Burma, having ∼3 and ∼8 times higher mean *in-situ* speciation events (mean: 64.04) and ∼3 times more subset sympatric speciation events (mean: 2.13) as compared to PI, with Wallacea and Sahul having very low sympatric speciation events (Table S6). Within PI and Wallacea and Sahul, however, *in-situ* speciation was followed by range-expansion and founder-event speciation, respectively (Table S6). Indo-Burma acted as a source region, with most of the emigration events occurring from Indo-Burma to PI (B→A; mean: 1.78 and 0.83, range expansion and founder-event speciation respectively) and from Indo-Burma to Wallacea and Sahul (B→C; mean: 2.34 and 1.59, range expansion and founder-event speciation respectively) during the Late Miocene (∼9‒7 Myr; Fig 3). Temporal examination of *in-situ* speciation rates showed that it peaked in Indo-Burma at ∼10‒9 Myr and again at 6 Myr (Figure 3B), in PI, it peaked at ∼6 Myr (Fig 3A), and in Wallacea and Sahul, it peaked during the Pliocene-Pleistocene boundary at ∼3‒2 Myr (Fig 3C).

**Fig 3.**
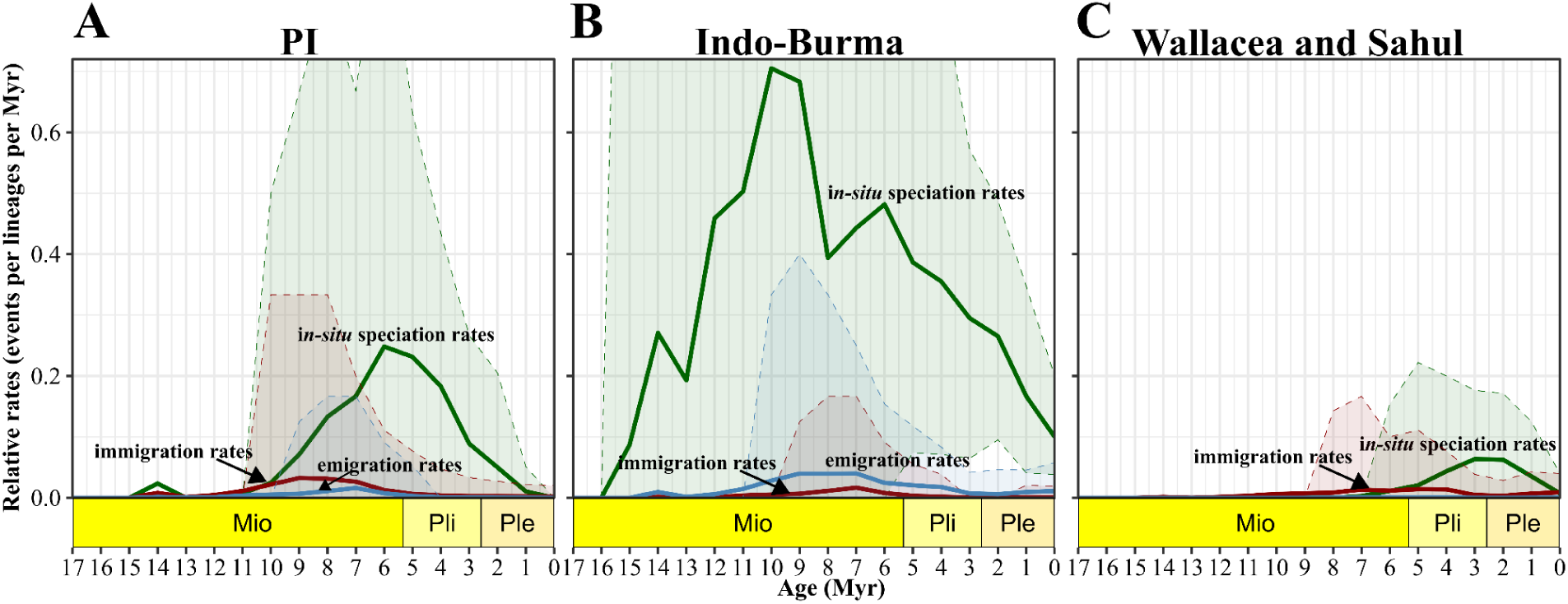
The relative rates of *in-situ* speciation, emigration, and immigration per 1 Myr time bins summarised across 100 posterior trees and 20 biogeographic stochastic mappings (BSMs) for each of the three biogeographic regions. Darker lines indicate the median rates, while the shaded ribbons indicate the lower 2.5% and upper 97.5% quantiles. Note that in panels A and B, the y-axis is clipped to ease the visual comparison. Green, blue and red lines indicate *in-situ* speciation rates, emigration rates and immigration rates, respectively. Mio: Miocene, Pli: Pliocene, Ple: Pleistocene.

### 3.4 Geographic variation in speciation rates across biogeographic subregions

The LTST plots indicate that lineage accumulation began early in Indo-Burma and increased rapidly after 10 Myr (Fig. 4A). Around the same time, lineages in PI also began accumulating, though the PI lineages showed signs of saturation from the Pleistocene onwards (Fig. 4A). In contrast, lineage accumulation in Wallacea and Sahul, however, began only after 5 Myr (Fig. 4A). Comparisons of tip-based speciation rates using ClaDS showed that Wallacea and Sahul, and Indo-Burma exhibited comparable speciation rates (mean: 0.21), while PI had a lower average speciation rate of 0.18 (Fig 4B; Table S6). Speciation rate estimates from DR also showed similar trends (Table S6).

**Fig 4.**
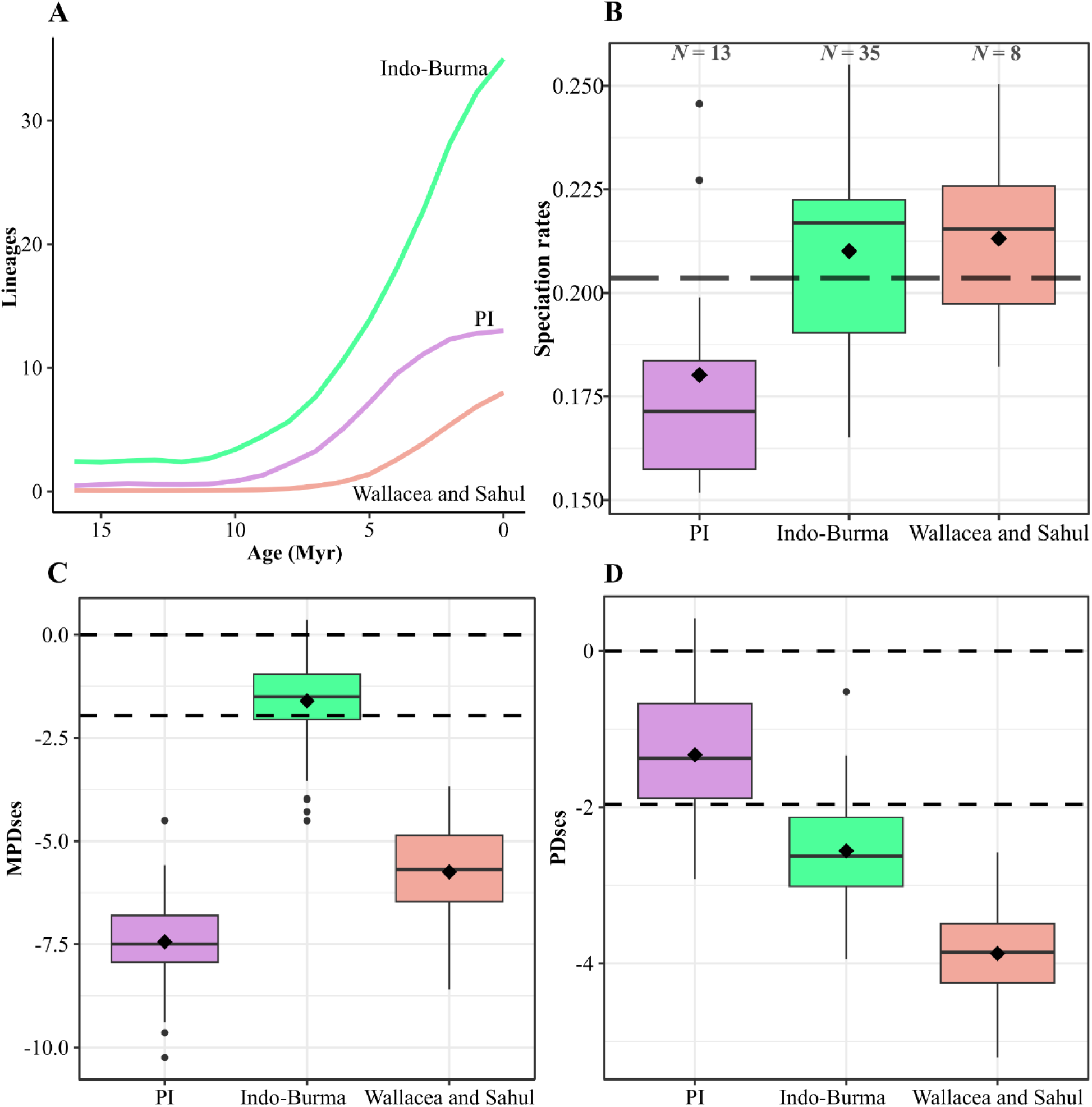
**A**. Mean lineage accumulation through time (1 Myr time bins) across 100 trees (sampled from the posterior) for each of the three regions. **B**. Tip-based speciation rates for each region estimated using ClaDS across 100 trees. The dashed line indicates the average tip rate, summarised across the entire *Miliusa* for 100 trees. **C** and **D**, standardised effect sizes of mean pairwise distance (MPDses) and phylogenetic diversity (PDses), respectively for each region, summarised over 100 trees. Dashed lines at 0 and -1.96 to indicate if the assemblage is random or clustered. The box plots represent the median and the quartile with the whiskers indicating 1.5 times the interquartile range. Diamonds represent the mean values for each region. The number of species per region is indicated by *N*.

### 3.5 Phylogenetic assemblage structure across biogeographic subregions

The PD of *Miliusa* lineages across the three regions mirrored the species richness patterns, with Indo-Burma having approximately twice the PD of PI and four times the PD of Wallacea and Sahul (Table S6). In terms of MPD, assemblages in PI and Wallacea and Sahul had similar values (mean: ∼15 and ∼14 Myr, respectively), while the MPD for assemblage in Indo-Burma was higher (∼20 Myr). MPDses, which is sensitive to deeper phylogenetic levels, indicated a random assemblage in Indo-Burma, while both PI and Wallacea and Sahul showed significant clustering (Fig. 4C). In contrast, PDses, which is more sensitive to the terminal level of the phylogeny, showed a random assemblage in PI, but strong clustering in Indo-Burma and Wallacea and Sahul (Fig 4D).

### 3.6 Climatic niche space occupied by the endemic PI species

In PI, most *Miliusa* species are geographically confined to the central and southern Western Ghats, except for four species: two endemic *M. sp1* and *M. sp2,* which occur in the drier Eastern Ghats, and *M. tomentosa* and *M. velutina*, which are widely distributed across PI and Indo-Burma (Fig 5). The endemic radiation in PI consisted of eleven species across the Western and the Eastern Ghats (node: 93, Fig. 2; Fig. 5). Of the nine species restricted to the Western Ghats, five occur south of the Palghat Gap, a major biogeographic barrier (below 10°N latitude), in the southern Western Ghats (Fig. 5). Despite their high geographical proximity, these species exhibit niche differentiation along climatic axes of water availability (precipitation seasonality and annual precipitation) and elevation (Fig 5; Table S7).

**Fig 5.**
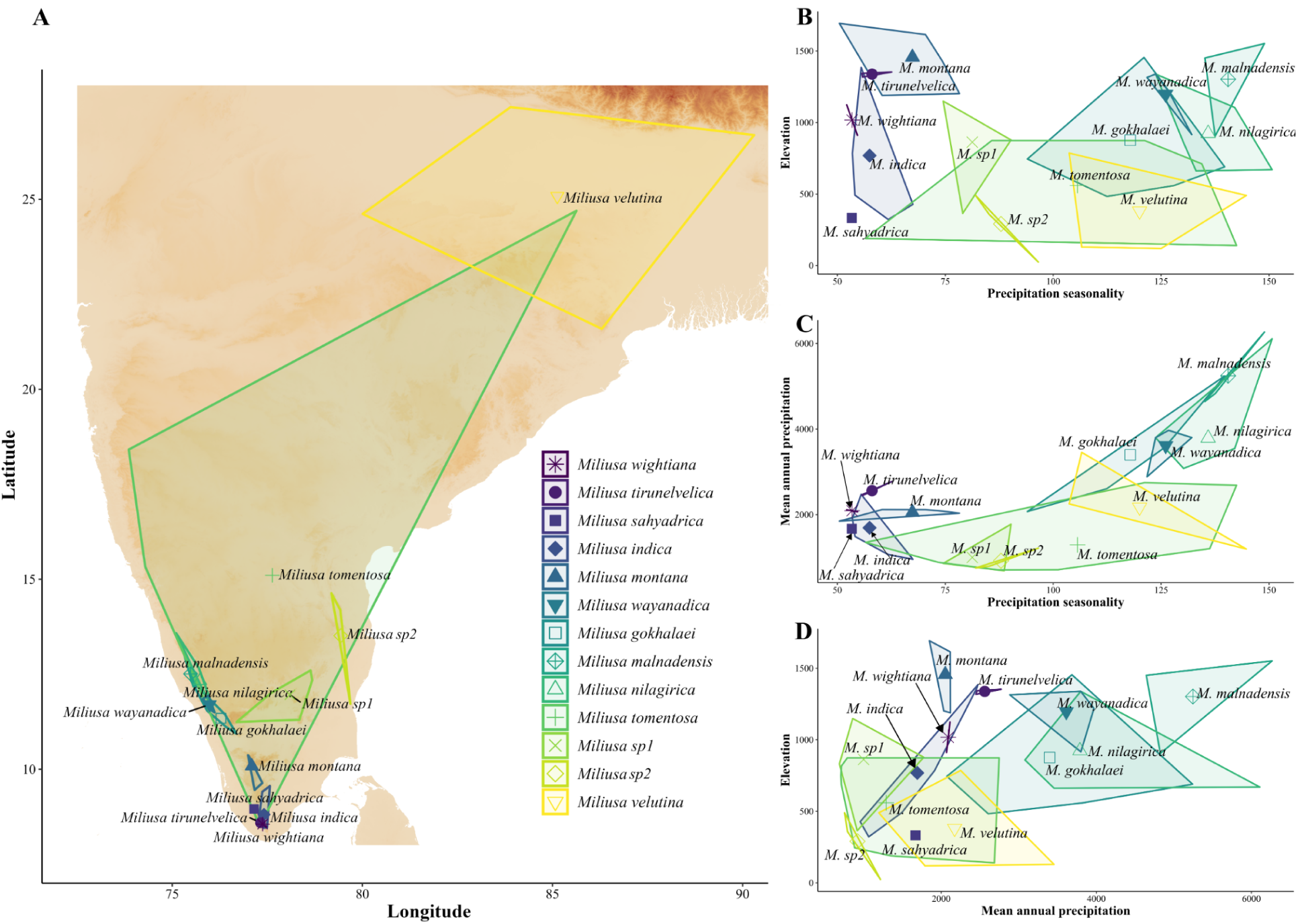
**A.** Geographical extent of *Miliusa* species found in peninsular India, plotted as a convex hull. Note that *M. tomentosa* and *M. velutina* have a wider distribution extending into Indo-Burma, which are not shown here. **B‒D**, climatic niche space occupied by each of the 13 species along the axes of precipitation seasonality, mean annual precipitation, and elevation, plotted as a convex hull. Symbols at the centroid of the hulls indicate the average value for each species along the respective climatic axes (niche position).

## 4. Discussion

Our results indicate that *Miliusa* originated in the mid-Miocene in Indo-Burma and PI. While most of the extant diversity is attributed to *in-situ* speciation, key dispersal events from Indo-Burma, early in the evolutionary history of *Miliusa*, likely spurred radiation in PI and Wallacea and Sahul. Although the rates of *in-situ* speciation events were higher and occurred earlier in Indo-Burma and PI as compared to Wallacea and Sahul, the speciation rates of lineages were the highest in Wallacea and Sahul, comparable to Indo-Burma, and were the lowest in PI. These biogeographic and diversification dynamics are reflected in the phylogenetically clustered assembly across regions, indicating the role of *in-situ* diversification in shaping the assemblages. Taken together, our results suggest that the observed asymmetry in extant diversity across regions is shaped by the interplay of time available for speciation, variation in speciation rates and dispersal dynamics, although their relative influence varied across regions. These results reinforce the existing body of evidence highlighting the crucial role of Indo-Burma in shaping tropical Asian biogeography and extant diversity, and emphasise the importance of Miocene and Pliocene-Pleistocene climatic fluctuations in driving speciation patterns.

### 4.1. Historical biogeography of *Miliusa*

At the deepest phylogenetic level, *Miliusa* comprises two sister clades. One clade being restricted to PI and Indo-Burma (node: 77; Clade A and B; Fig. 2), while the other is primarily an Indo-Burman clade (node: 54; Clade C and D; Fig. 2) with subsequent founder-event speciation resulting in the establishment of lineages in Wallacea and Sahul during the Pliocene (node: 73; Fig. 2). The PI and Indo-Burman clade (Clade A and B), is characterised by subset sympatric speciation events early in its evolutionary history, with vicariance, subset sympatry and range-expansion events shaping the lineages found in the PI. In contrast, all Indo-Burman lineages in this clade underwent *in-situ* speciation (node: 80; Fig. 2). For both Clade A and Clade B, *in-situ* speciation events predominantly occurred during the Pliocene. The sister clade (Clade C and D; Fig. 2) is largely characterised by *in-situ* speciation events during the Pliocene and Pleistocene, along with two range expansions and two founder-event speciation events from Indo-Burma to Wallacea and Sahul.

Indo-Burma is primarily characterised by *in-situ* speciation events, which were ∼3 and ∼8 times higher (mean: 64.04) as compared to PI and Wallacea and Sahul, and peaked during the late Miocene and at the Miocene–Pliocene boundary (Fig 3B). While overall dispersal events were low in *Miliusa*, most were from Indo-Burma to PI and Wallacea and Sahul, peaking during the Late Miocene (∼9‒7 Myr). The relative geo-climatic stability, large area and heterogeneous habitat of Indo-Burma likely supported the steady accumulation of *Miliusa* lineages throughout its evolutionary history. Our results support the role of Indo-Burma as an evolutionary hotspot for *Miliusa* facilitating lineage persistence, diversification and dispersal to other biogeographic regions, consistent with patterns observed for several other floral and faunal clades (Crayn et al., 2014; De Bruyn et al., 2014; Joyce et al., 2020).

In the case of PI, most of the cladogenetic and anagenetic events occurred during the Late Miocene and Early Pliocene, in congruence with the Miocene aridification and the intensification of the monsoon in the region (Ding et al., 2020; Morley et al., 2018). These climatic shifts resulted in the contraction of wet forests, which likely led to the climatic vicariance of lineages that were previously widespread across Indo-Burma and PI (node: 77, 78, 79, and 92). These results are consistent with patterns observed in other Annonaceae genera in the region, such as *Artabotrys* (Chen et al., 2019), *Goniothalamus* (Thomas et al., 2017), and *Meiogyne* (Thomas et al., 2012), all of which show evidence of climatic vicariance from a larger ancestral range of Indo-Burma and PI and dispersal from Indo-Burma to PI during the Late Miocene and Early Pliocene. The ancestral node (node: 93) of the endemic PI radiation is likely to have resulted from either a founder-event speciation (DEC+*j* model, log-likelihood: -35.93) or vicariance (DEC model log-likelihood: -37.18). Given the similar likelihood and AICc values of the two models, we hypothesise that both events are plausible given the geoclimatic and biogeographic context. Previous syntheses of biogeography patterns of several Annonacea clades in tropical Asia, with limited sampling from PI, show that PI lineages are often nested within Asian clades (Thomas et al., 2015). However, by including all known PI species of *Miliusa*, our results indicate that PI may have played a more prominent role in the early evolutionary history of the genus. We recovered an endemic radiation of 11 species in PI, and PI along with Indo-Burma, were recovered as the ancestral area for *Miliusa.* Ancestral range estimations across multiple biogeographic histories also recovered a low frequency of dispersal events from PI to Indo-Burma (Fig 3A), indicating its limited role in shaping the regional assembly from the end Miocene onwards.

We found that the main radiation of *Miliusa* in Wallacea and Sahul likely resulted from founder-event speciation or vicariance from Indo-Burma (node: 72) during the Late Miocene-Pliocene, followed by *in-situ* speciation. In addition, two range expansion events, expanding ranges of *M. horsfieldii and M. cf horsfieldii* and one founder-event speciation from Indo-Burma leading to *M. butonensis* during the Pliocene-Pleistocene were also detected, with the rates of *in-situ* speciation in Wallacea and Sahul peaking at the Pliocene-Pleistocene boundary (Fig 3C). These results add *Miliusa* to the list of plant lineages that have successfully dispersed across Wallace’s Line from Indo-Burma to Wallacea and Sahul since the Late Miocene, followed by rapid speciation post-dispersal (Crayn et al., 2014; De Bruyn et al., 2014; Joyce et al., 2020; Zhang et al., 2022). These patterns were likely facilitated by the reduced distance between the landmasses, favourable climatic conditions and the emergence of new niches due to the upliftment of landmasses in Wallacea and parts of Sahul, allowing for both dispersal and *in-situ* speciation of lineages (De Bruyn et al., 2014; Kooyman et al., 2019; Lohman et al., 2011).

### 4.2. Geography of speciation rates, phylogenetic assembly, and diversity patterns

Indo-Burma, currently the centre of *Miliusa* diversity, consistently exhibited higher lineage diversity compared to other regions throughout the evolutionary history of the genus. Among the three regions, it exhibited the highest speciation rates and had ∼2 and ∼4 times higher PD than PI and Wallacea and Sahul, respectively. Indo-Burma’s relative climatic stability, large area and more heterogeneous habitats likely allowed for higher speciation rates due to potentially higher availability of niches (De Bruyn et al., 2014; Lohman et al., 2011). Its geological and climatic stability, especially during the Pliocene-Pleistocene, allowed for the persistence of wet rainforest lineages by acting as refugia during climatic perturbations (Lohman et al., 2011). Together, these features facilitated continuous lineage divergence over time and dispersal to other regions, reinforcing the role of Indo-Burma as an evolutionary hotspot. The biogeographic history and the speciation trends of *Miliusa* are reflected in the phylogenetic assemblage structure, which appears random at the basal level and strongly clustered at the terminal level. The random structure at the basal level is likely due to how the Indo-Burma assemblage has species from different subclades in the *Miliusa* phylogeny (nodes: 80, 65, and 55), resulting in the assemblage being composed of distantly related species at the basal level. In contrast, the strong clustering at the terminal level is indicative of the assemblage being composed of more closely related species due to *in-situ* speciation within each of these subclades.

In contrast to the other two biogeographic regions, lineages in PI, despite being as old as those in Indo-Burma, show much lower lineage diversity, signs of saturation, and markedly lower speciation rates (Fig. 4A, B). While PI has remained geologically stable post-collision with Asia, the onset of Miocene aridification resulted in the contraction of the wet forest to the wetter slopes of the Western Ghats and isolated high-elevation pockets in the Eastern Ghats (Morely et al., 2018). This reduction in key habitat for predominantly wet-affiliated *Miliusa* likely constrained the available niche for speciation in PI, with early-diverging lineages filling up most of the available niches. While *M. sp1*, *M. sp2,* and *M. indica* have expanded into relatively drier habitats and exhibit brevi-deciduous leaf phenology, the majority of the endemic species remain wet-affiliated. Of the eleven endemic species in PI, more than half are range-restricted. Our qualitative examination of climatic niche space also indicates niche differentiation along the gradients of water availability and elevation (Fig. 5). We hypothesise that niche conservatism for wet habitats, a pattern widely reported in many other Annonaceae clades (Couvreur et al., 2011; Thomas et al., 2015), might be a potential process driving speciation patterns of *Miliusa* in PI and possibly within Indo-Burma and Wallacea and Sahul as well, which needs to be examined further. In terms of the phylogenetic assemblage structure, PI lineages show a contrasting trend of strong clustering at the basal level (Fig. 4C), likely due to the assemblage being predominantly composed of species from a single subclade (node: 93). In contrast, there was no structure at the terminal level (Fig. 4D), which is likely due to distinct phylogenetic positions of *M. tomentosa* and *M. velutina* from Clade A, and *M. nilagirica,* in relation to the species in the endemic PI clade.

Lastly, lineages in Wallacea and Sahul, despite their relatively recent origin and with most *in-situ* speciation events occurring in the Pliocene, show rapid lineage accumulation and high speciation rates, consistent with island-like radiations. They exhibit speciation rates comparable to those of Indo-Burma and 1.2 times higher median speciation rates than lineages in PI, despite having the lowest PD and species richness among the three regions. These patterns also influence the phylogenetic assemblage structure, which shows strong clustering at both the basal and terminal levels of the phylogeny, indicating that the assemblage is composed of closely related species at different phylogenetic depths. Tectonic movements, recurrent climatic oscillations and upliftment of landmasses in the region likely opened up new niches for speciation and allowed for dispersal-driven radiation in this region (Crayn et al., 2014; De Bruyn et al., 2014; Joyce et al., 2020).

Collectively, we find that the spatial and temporal disparities in *Miliusa* diversity are shaped by regional attributes, primarily geo-climatic stability and niche availability, which allow clades to diversify over longer periods (as in Indo-Burma), at faster rates (as in Wallacea and Sahul), or limit speciation (as in PI). Within-region *in-situ* speciation events are the primary determinants of *Miliusa* diversity, with limited influence of dispersal. However, dispersal (or vicariance) played a pivotal role in spurring the diversification of lineages in both PI and Wallacea and Sahul. Overall, the interaction between regional characteristics and key macroevolutionary processes shape the regional diversity patterns of *Miliusa.* However, the relative importance of each of these processes varies with regional geo-climatic and biogeographic context.

### 4.4 Conclusion

The variation in geo-climatic stability and niche availability across regions, along with the biogeographic and diversification dynamics, shaped the asymmetric diversity patterns of *Miliusa* within tropical Asia. *In-situ* speciation is the primary determinant of diversity in the region, with Indo-Burma being the centre for diversity for *Miliusa* and an evolutionary hotspot, allowing for both the persistence and diversification of lineages. Meanwhile, Wallacea and Sahul exhibit patterns typical of island radiation, characterised by rapid, recent lineage accumulation post-dispersal. The inclusion of all the PI lineages indicates that PI likely played a significant role in the early evolutionary history of *Milisua,* although its overall influence on broader diversity patterns across tropical Asia appears limited. Collectively, our results emphasise the role of Miocene-driven aridification and Pliocene-Pleistocene climatic fluctuations in shaping the diversification dynamics of *Miliusa* across tropical Asia.

## Acknowledgements

This study was funded by DST-SERB as part of the National Post Doctoral Fellowship (NPDF: Grant number: PDF/2016/000865) awarded to NP. NP would like to thank the Director of CSIR-Central Institute of Medicinal and Aromatic Plants (CSIR-CIMAP) for hosting him as an NPDF at the Bengaluru Research Centre. We would like to thank the respective forest departments of the states of Andaman and Nicobar, Andhra Pradesh, Arunachal Pradesh, Assam, Karnataka, Kerala, Maharashtra, Tamil Nadu, and Uttarakhand for granting us the research permits for this work. NP thanks to Dr Gokul and Dr K Ravikumar for facilitating the field work. We would like to extend our sincere thanks to Dr. Praveen Karanth (Indian Institute of Sciences, Bengaluru) for providing laboratory space and access to PCR facilities, which were used to amplify some of the markers included in the study. AG was supported by a start-up grant awarded to J.J. from CSIR-Centre for Cellular and Molecular Biology (CSIR-CCMB), Uppal Road, Hyderabad, India. We thank the DNA sequencing facility and the IT team for access to the HPC facility at the CSIR-CCMB. AG thanks Pragyadeep Roy, Rishiddh Jhaveri, Hannah Krupa, and members of the Evol-Eco lab for discussions and inputs. AG thanks Pragyadeep Roy for the discussion and inputs on the analyses and for sharing the scripts for the speciation analyses. AG thanks Hannah Krupa for proofreading the manuscript. AG thanks Dr. Bharti Dharapuram, Kaikho Liriina, Dr. Mihir Kulkarni, and Nehal Gurung for their help and inputs in the lab work.

## Conflict of Interest

The authors declare no conflict of interest.

## Data Availability

The molecular data generated in this study will be deposited in NCBI. Additional details are provided in the supplementary.

## Author Contributions

NP designed the study and conducted the fieldwork. AK, NT, and AG generated the molecular data. AG, JJ, and NP conceptualised the manuscript. AG led the analysis and writing of the manuscript with inputs from JJ and NP. VS supported the project from the inception stage and facilitated research permits and field work. All the authors reviewed and edited the manuscript.

## Notes

### Competing Interest Statement

The authors have declared no competing interest.

