## Supplementary for "*In-situ* diversification and regional attributes shape asymmetric diversity of *Miliusa* (Annonaceae) in tropical Asia"

Table S1. List of *Miliusa* species as recognised by the KEW Plants of the world, and their distribution is shown. The distribution column indicates the biogeographic distribution as used in this study. Undescribed species, which are morphologically and phylogenetically distinct, are indicated by grey cells. Species treated as synonyms are reported in rows 69-73.

| Sno | Full name authority | Distribution | DNA Sequence Data |
|---|---|---|---|
| 1 | *Miliusa amplexicaulis* Ridl. | Indo-Burma | Downloaded from GenBank |
| 2 | *Miliusa balansae* Finet & Gagnep. | Indo-Burma | Downloaded from GenBank |
| 3 | *Miliusa brahei* (F.Muell.) Jessup | Wallacea and Sahul | Downloaded from GenBank |
| 4 | *Miliusa butonensis* Chaowasku & Kessler | Wallacea and Sahul | Downloaded from GenBank |
| 5 | *Miliusa campanulata* Pierre | Indo-Burma | Downloaded from GenBank |
| 6 | *Miliusa chantaburiana* Damth. & Chaowasku | Indo-Burma | Downloaded from GenBank |
| 7 | *Miliusa cuneata* Craib | Indo-Burma | Downloaded from GenBank |
| 8 | *Miliusa eupoda* (Miq.) I.M.Turner | Indo-Burma | Downloaded from GenBank |
| 9 | *Miliusa fragrans* Chaowasku & Kessler | Indo-Burma | Downloaded from GenBank |
| 10 | *Miliusa fusca* Pierre | Indo-Burma | Downloaded from GenBank |
| 11 | *Miliusa glandulifera* C.E.C.Fisch. | Indo-Burma | Downloaded from GenBank |
| 12 | *Miliusa horsfieldii* (Benn.) Baill. ex Pierre | Indo-Burma and Wallacea and Sahul | Downloaded from GenBank |
| 13 | *Miliusa cf horsfieldii* | Indo-Burma and Wallacea and Sahul | Downloaded from GenBank |
| 14 | *Miliusa intermedia* Chaowasku & Kessler | Indo-Burma | Downloaded from GenBank |
| 15 | *Miliusa koolsii* (Kosterm.) J.Sinclair | Wallacea and Sahul | Downloaded from GenBank |
| 16 | *Miliusa lanceolata* Chaowasku & Kessler | Wallacea and Sahul | Downloaded from GenBank |
| 17 | *Miliusa macrocarpa* Hook.f. & Thomson | Indo-Burma | Downloaded from GenBank |
| 18 | *Miliusa macropoda* Miq. | Indo-Burma | Downloaded from GenBank |
| 19 | *Miliusa majestatis* Damth., Sinbumr. & Chaowasku | Indo-Burma | Downloaded from GenBank |
| 20 | *Miliusa microphylla* Damth. & Chaowasku | Indo-Burma | Downloaded from GenBank |

| | | | |
|---|---|---|---|
| 21 | *Miliusa mollis* Pierre | Indo-Burma | Downloaded from GenBank |
| 22 | *Miliusa nakhonsiana* Chaowasku & Kessler | Indo-Burma | Downloaded from GenBank |
| 23 | *Miliusa novoguineensis* Mols & Kessler | Wallacea and Sahul | Downloaded from GenBank |
| 24 | *Miliusa parviflora* Ridl. | Indo-Burma | Downloaded from GenBank |
| 25 | *Miliusa pumila* Chaowasku | Indo-Burma | Downloaded from GenBank |
| 26 | Miliusa sclerocarpa (A.DC.) Kurz | Indo-Burma | Downloaded from GenBank |
| 27 | *Miliusa sessilis* Chaowasku & Kessler | Indo-Burma | Downloaded from GenBank |
| 28 | *Miliusa sp2 Thailand* | Indo-Burma | Downloaded from GenBank |
| 29 | *Miliusa thailandica* Chaowasku & Kessler | Indo-Burma | Downloaded from GenBank |
| 30 | *Miliusa thorelii* Finet & Gagnep. | Indo-Burma | Downloaded from GenBank |
| 31 | *Miliusa traceyi* Jessup | Wallacea and Sahul | Downloaded from GenBank |
| 32 | *Miliusa umpangensis* Chaowasku & Kessler | Indo-Burma | Downloaded from GenBank |
| 33 | *Miliusa andamanica* (King) Finet & Gagnep. | Indo-Burma | Generated in this study |
| 34 | *Miliusa dioeca* (Roxb.) Chaowasku & Kessler | Indo-Burma | Generated in this study |
| 35 | *Miliusa gokhalaei* Ratheesh, Sujanapal, Anil Kumar & Sivad. | PI | Generated in this study |
| 36 | *Miliusa cf horsfieldii* Andaman | Indo-Burma | Generated in this study |
| 37 | *Miliusa indica* Lesch. ex A.DC. | PI | Generated in this study |
| 38 | *Miliusa malnadensis* N.V.Page & Nerlekar | PI | Generated in this study |
| 39 | *Miliusa cf montana* Gardner ex Hook.f. & Thomson | PI | Generated in this study |
| 40 | *Miliusa nilagirica* Bedd. | PI | Generated in this study |
| 41 | *Miliusa cf parviflora Andaman* | Indo-Burma | Generated in this study |
| 42 | *Miliusa sahyadrica* G.Rajkumar, Alister, Nazarudeen & Pandur. | PI | Generated in this study |
| 43 | *Miliusa sp 1* | PI | Generated in this study |
| 44 | *Miliusa sp1 Andaman* | Indo-Burma | Generated in this study |

| | | | |
|---|---|---|---|
| 45 | *Miliusa sp2* | PI | Generated in this study |
| 46 | *Miliusa sp2 Andaman* | Indo-Burma | Generated in this study |
| 47 | *Miliusa sp3 Andaman* | Indo-Burma | Generated in this study |
| 48 | *Miliusa tirunelvelica* Murugan, Manickam, Sundaresan & Jothi | PI | Generated in this study |
| 49 | *Miliusa tomentosa* (Roxb.) Finet & Gagnep. | PI and Indo-Burma | Generated in this study |
| 50 | *Miliusa velutina* (DC.) Hook.f. & Thomson | PI and Indo-Burma | Generated in this study and downloaded from NCBI |
| 51 | *Miliusa wayanadica* Sujanapal, Ratheesh & Sasidh. | PI | Generated in this study |
| 52 | *Miliusa wightian*a Hook.f. & Thomson | PI | Generated in this study |
| 53 | *Miliusa astiana* Chaowasku & Kessler | Indo-Burma | Not represented in this study |
| 54 | *Miliusa baillonii* Pierre | Indo-Burma | Not represented in this study |
| 55 | *Miliusa banghoiensis* Jovet-Ast | Indo-Burma | Not represented in this study |
| 56 | *Miliusa cambodgensis* Chaowasku & Kessler | Indo-Burma | Not represented in this study |
| 57 | *Miliusa caudata* N.Balach. & Chakrab. | Indo-Burma | Not represented in this study |
| 58 | *Miliusa codonantha* Chaowasku | Indo-Burma | Not represented in this study |
| 59 | *Miliusa filipes* Ridl. | Indo-Burma | Not represented in this study |
| 60 | *Miliusa glochidioides* Hand.-Mazz | Indo-Burma | Not represented in this study |
| 61 | *Miliusa hirsuta* Chaowasku & Kessler | Indo-Burma | Not represented in this study |
| 62 | *Miliusa ninhbinhensis* Chaowasku & Kessler | Indo-Burma | Not represented in this study |
| 63 | *Miliusa parvifolia* (Kurz) Damth. & Chaowasku | Indo-Burma | Not represented in this study |
| 64 | *Miliusa saccata* C.E.C.Fisch. | Indo-Burma | Not represented in this study |
| 65 | *Miliusa tenuistipitata* W.T.Wang | Indo-Burma | Not represented in this study |
| 66 | *Miliusa tristi*s Kurz | Indo-Burma | Not represented in this study |
| 67 | *Miliusa vidalii* J.Sinclair | Indo-Burma | Not represented in this study |
| 68 | *Miliusa viridiflora* Chaowasku & Kessler | Wallacea and Sahul | Not represented in this study |

| | | | |
|---|---|---|---|
| 69 | *Miliusa manickamiana* Murugan | PI | Treated as a synonym of *M. indica* |
| 70 | *Miliusa paithalmalayana* Josekutty | PI | Treated as a synonym of *M. malnadensis* |
| 71 | *Miliusa agasthyamalana* V.S.A.Kumar & Sindhu Arya | PI | Treated as a synonym of *M. tirunelvelica* |
| 72 | *Miliusa ammaiae* Karupp. & P.S.S.Rich. | PI | Treated as a synonym of *M. wightiana* |
| 73 | *Miliusa flaviviridis* N.V.Page, Poti & K.Ravik. | PI | Treated as a synonym of *M. wightiana* |

Table S2. Details of the molecular data generated in the study and the GenBank accession numbers for sequences used for phylogenetic reconstruction are shown. — indicates no sequence. For the primary data, ndhF was split into two non-overlapping fragments ndhF1 and ndhF2.

| **Sno** | **Species** | **Location** | **trnLF** | **matK** | **ndhf1** | **ndhf2** | **psbA-trnH** | **ycf1** |
|---|---|---|---|---|---|---|---|---|
| **Primary data** | | | | | | | | |
| 1 | *Miliusa andamanica* | Andaman, Havelock | Y | Y | Y | Y | Y | Y |
| 2 | *Miliusa cf horsfieldii Andaman* | Andaman, Little Andaman, Hut Bay | Y | Y | — | — | Y | Y |
| 3 | *Miliusa cf montana ind 1* | Tamil Nadu, Megmalai Wildlife Sanctuary | Y | Y | Y | Y | Y | Y |
| 4 | *Miliusa cf montana ind 2* | Kerala, Periyar Tiger Rerve, Pandantoda | Y | Y | Y | Y | Y | Y |
| 5 | *Miliusa cf montana ind 3* | Tamil Nadu, Valparai, Akkamalai | Y | Y | — | Y | Y | Y |
| 6 | *Miliusa cf parviflora Andaman ind 1* | Andaman, Little Andaman, HutBay | Y | Y | Y | Y | Y | Y |
| 7 | *Miliusa cf parviflora Andaman ind 2* | Andaman, Little Andaman, HutBay | Y | Y | — | — | Y | Y |
| 8 | *Miliusa dioeca ind 1* | Assam, Garampani Wildlife Sanctuary | Y | Y | Y | Y | Y | Y |
| 9 | *Miliusa dioeca ind 2* | Arunachal Pradesh, Pakke | — | Y | — | — | Y | Y |
| 10 | *Miliusa gokhalaei* | Kerala, Nilambur, Nadukani Ghat | Y | Y | — | Y | Y | Y |
| 11 | *Miliusa indica ind1* | Tamil Nadu, Kalakad Mundunthurai Tiger Reserve | Y | Y | — | Y | Y | Y |
| 12 | *Miliusa indica ind 2* | Tamil Nadu, Kalakad Mundunthurai Tiger Reserve | Y | Y | — | — | Y | Y |
| 13 | *Miliusa indica ind 3* | Tamil Nadu, Sriviliputtur | Y | Y | Y | Y | Y | — |
| 14 | *Miliusa malnadensis* | Karnataka, Kudremukh peak | Y | — | — | Y | — | |
| 15 | *Miliusa nilagirica ind 1* | Kerala, Wayanad, 900 forest | Y | Y | Y | Y | Y | Y |

| | | | | | | | | |
|---|---|---|---|---|---|---|---|---|
| 16 | *Miliusa nilagirica ind 2* | Karnataka, Agumbe, Kalinga estate | Y | Y | Y | Y | Y | Y |
| 17 | *Miliusa sahyadrica* | Kerala, Palaruvi | — | Y | Y | Y | Y | Y |
| 18 | *Miliusa sp1* | Tamil Nadu, MM hills | Y | Y | Y | Y | Y | — |
| 19 | *Miliusa sp1 Andaman ind 1* | South Andaman, Chidiatapu | Y | Y | — | Y | Y | Y |
| 20 | *Miliusa sp1 Andaman ind 2* | South Andaman, Chidiatapu | — | Y | — | — | Y | Y |
| 21 | *Miliusa sp2* | Tamil Nadu, Pondicherry | Y | Y | Y | Y | Y | Y |
| 22 | *Miliusa sp2* | Andhra Pradesh, Chitvel | Y | Y | Y | Y | Y | Y |
| 23 | *Miliusa sp2 Andaman* | Andaman, Rutland | Y | Y | Y | Y | Y | Y |
| 24 | *Miliusa sp3 Andaman* | Andaman, Havelock, Elephant beach | Y | Y | — | Y | Y | Y |
| 25 | *Miliusa tirunelvelica ind 1* | Tamil Nadu, Kalakad Mundunthurai Tiger Reserve, Kakachi | Y | Y | Y | Y | Y | Y |
| 26 | *Miliusa tirunelvelica ind 2* | Tamil Nadu, Kalakad Mundunthurai Tiger Reserve, Kakachi | Y | Y | Y | Y | Y | Y |
| 27 | *Miliusa tomentosa ind 1* | Karnataka, Bangalore, Valley School | Y | Y | Y | Y | Y | Y |
| 28 | *Miliusa tomentosa ind 2* | Pune, Katraj | Y | Y | Y | Y | Y | Y |
| 29 | *Miliusa velutina ind 1* | Uttarakhand, Rajaji | Y | Y | — | Y | Y | Y |
| 30 | *Miliusa wayanadica* | Kerala, Wayanad, Periya Reserve Forest | Y | Y | Y | Y | Y | Y |
| 31 | *Miliusa wightiana* | Tamil Nadu, Kalakad Mundunthurai Tiger Reserve, Naraikadu | Y | Y | — | — | Y | — |
| **Data downloaded from NCBI** | | | | | | | | |
| | | | **trnLF** | **matK** | **ndhF** | | **psbA-trnH** | **ycf1** |
| 32 | *Miliusa amplexicaulis* | | JQ690478 | — | JQ690479 | | JQ690480 | JQ690481 |
| 33 | *Miliusa balansae* | | JQ690482 | — | JQ690483 | | JQ690484 | JQ690485 |
| 34 | *Miliusa brahei* | | JQ690430 | — | JQ690431 | | JQ690432 | JQ690433 |
| 35 | *Miliusa butonensis* | | JQ690434 | — | JQ690435 | | JQ690436 | JQ690437 |
| 36 | *Miliusa campanulata* | | JQ690486 | — | JQ690487 | | JQ690488 | JQ690489 |
| 37 | *Miliusa chantaburiana* | | OL438845 | MH663444 | OL438791 | | OL438817 | OL438864 |
| 38 | *Miliusa cuneata* | | JQ690490 | — | JQ690491 | | JQ690492 | JQ690493 |
| 39 | *Miliusa eupoda* | | OL438847 | OL438773 | OL438793 | | OL438819 | OL438866 |
| 40 | *Miliusa fragrans* | | JQ690438 | — | JQ690439 | | JQ690440 | JQ690441 |
| 41 | *Miliusa fusca* | | JQ690442 | — | JQ690443 | | JQ690444 | JQ690445 |
| 42 | *Miliusa glandulifera* | | OL438859 | OL438784 | OL438809 | | OL438831 | OL438882 |
| 43 | *Miliusa horsfieldii* | | JQ690446 | — | JQ690447 | | JQ690448 | JQ690449 |
| 44 | *Miliusa cf horsfieldii* | | OL438861 | OL438785 | OL438811 | | OL438833 | OL438884 |
| 45 | *Miliusa intermedia* | | JQ690450 | — | JQ690451 | | JQ690452 | JQ690453 |
| 46 | *Miliusa koolsii* | | JQ690454 | — | JQ690455 | | JQ690456 | JQ690457 |

| | | | | | | | |
|---|---|---|---|---|---|---|---|
| 47 | *Miliusa lanceolata* | | JQ690458 | — | JQ690459 | JQ690460 | JQ690461 |
| 48 | *Miliusa macrocarpa* | | JQ690498 | JQ690499 | JQ690500 | JQ690501 | JQ690502 |
| 49 | *Miliusa macropoda* | | JQ690462 | — | JQ690463 | JQ690464 | JQ690465 |
| 50 | *Miliusa majestatis* | | OR267123 | OR267124 | OR267125 | OR267122 | OR267127 |
| 51 | *Miliusa microphylla* | | AY319102 | AY518851 | JQ690503 | JQ690504 | JQ690505 |
| 52 | *Miliusa mollis* | | OL438848 | OL438774 | OL438796 | OL438820 | OL438869 |
| 53 | *Miliusa nakhonsiana* | | OL438858 | OL438783 | OL438808 | OL438830 | OL438881 |
| 54 | *Miliusa novoguineensis* | | JQ690466 | — | JQ690467 | JQ690468 | JQ690469 |
| 55 | *Miliusa parviflora* | | JQ690470 | — | JQ690471 | JQ690472 | JQ690473 |
| 56 | *Miliusa pumila* | | JQ690511 | MH663447 | OL438794 | JQ690513 | OL438867 |
| 57 | *Miliusa pumila* | | JQ690474 | — | JQ690475 | JQ690476 | JQ690477 |
| 58 | *Miliusa sclerocarpa* | | JQ690474 | — | JQ690475 | JQ690476 | JQ690477 |
| 59 | *Miliusa sessilis* | | OL438855 | OL438781 | OL438805 | OL438827 | OL438878 |
| 60 | *Miliusa sp2 Thailand* | | JQ690526 | JQ690527 | JQ690528 | JQ690529 | JQ690530 |
| 61 | *Miliusa thailandica* | | JQ690515 | — | JQ690516 | JQ690517 | JQ690518 |
| 62 | *Miliusa thorelii* | | AY319104 | AY518846 | JQ690519 | JQ690520 | JQ690521 |
| 63 | *Miliusa traceyi* | | JQ690531 | JQ690532 | JQ690533 | JQ690534 | JQ690535 |
| 64 | *Miliusa umpangensis* | | JQ690522 | — | JQ690523 | JQ690524 | JQ690525 |
| 65 | *Miliusa velutina ind 2* | | AY319105 | AY518847 | JQ690536 | JQ690537 | JQ690538 |
| 66 | *Miliusa dioeca ind 3* | | JQ690494 | — | JQ690495 | JQ690496 | JQ690497 |

Table S3. Primer pairs and amplification conditions for PCRs used in the study. The primer selection and PCR protocols were done following Chaowasku et al., (2012). These markers have been extensively used for phylogenetic reconstruction for *Miliusa* and other genera in Annonaceae.

| | **Primer name** | **Primer sequences (5'-3')** | **Amplification conditions** |
|---|---|---|---|
| ***matK*** | 390 F | CGATCTATTCATTCAATATTTC | 94°C – 5 min; 94°C – 1 min, 48°C – 30 sec, 72°C – 1 min, 30 cycles; 72°C – 7 min |
| | 1326 R | TCTAGCACACGAAAGTCGAAGT | |
| ***ycf1*** | 914F | GGATGGGAATGAATGAAGAAATGC | 95°C – 5 min; 95 °C – 1 min, 50 °C – 1 min, 65 °C – 4 min, 40 cycle; 65°C – 5 min ) |
| | 2323R | CCGTATCAATATGCTTGTCCTC | |
| ***trnLF*** | trnl-c | CGAAATCGGTAGACGCTACG | 95°C – 5 min; 95 °C – 1 min, 50 °C – 1 min, 65 °C – 4 min, 40 cycle; 65°C – 5 min ) |
| | trnL-f | ATTTGAACTGGTGACACGA | |
| ***ndhF*** | 54 F | GCTCGTCGTATGTGGGCTTTT | 94°C – 4 min; 94 °C – 30 sec, 54 °C – 1 min, 72 °C – 2 min, 35 cycle; 72°C – 7 min |
| | 1650 R | CGAAGGGAATTCCTATGGACC | |
| ***psbA-trnH*** | psbA | CGAAGCTCCATCTACAAATGG | 94°C – 5 min; 94 °C – 1 min, 55 °C – 1 min, 72 °C – 1 min, 35 cycle; 72°C – 7 min |
| | trnH | ACTGCCTTGATCCACTTGGC | |

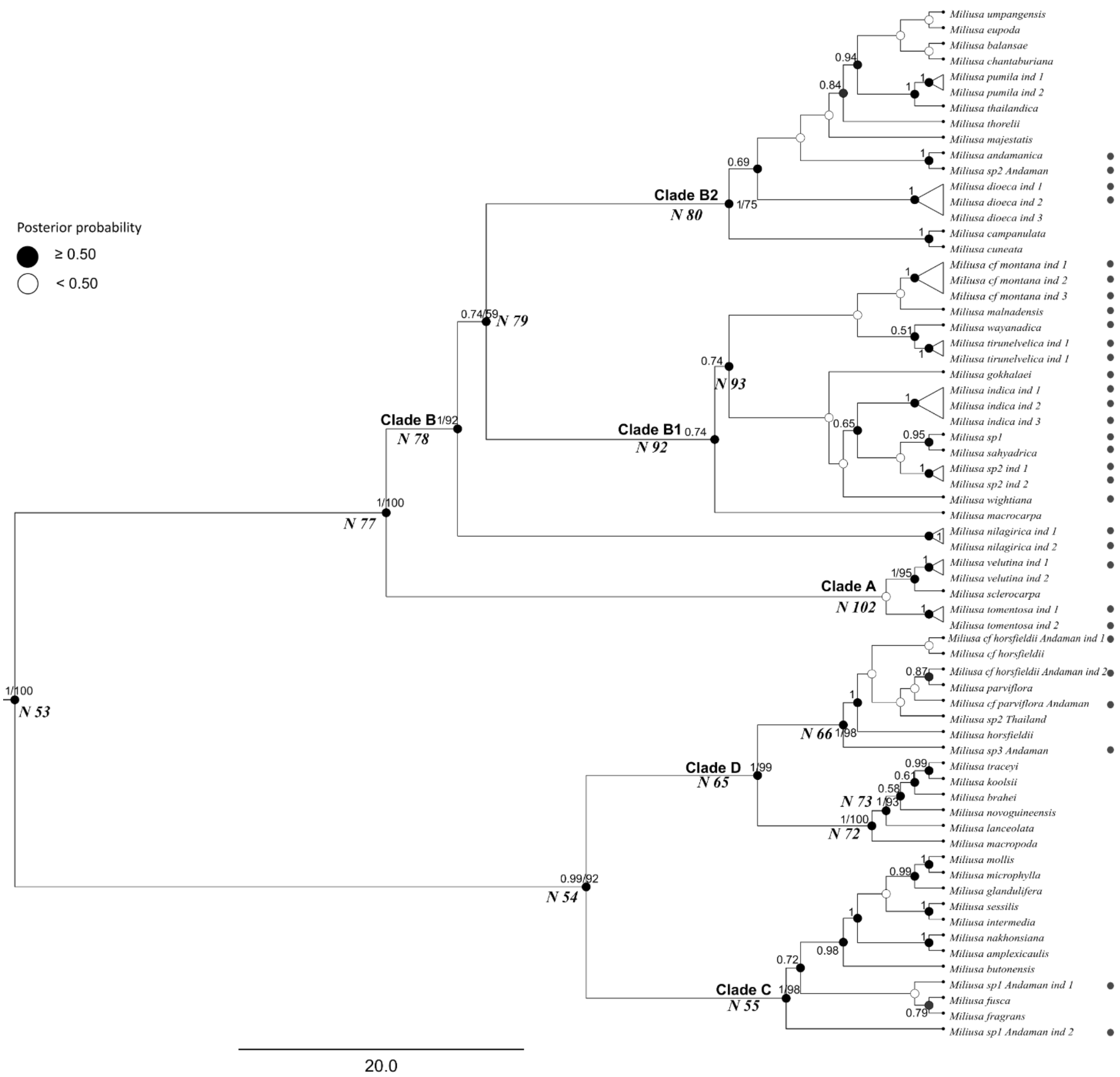


Fig S1: Bayesian phylogenetic tree based on the concatenated plastid data (five markers). Bayesian posterior probability values (PP) and maximum likelihood (ML) bootstrap values greater than 50% are indicated at each node (PP/ML). Select key clades and nodes are labelled and numbered. Circles next to the species name indicate the accessions for which molecular data was generated in the study. Please see the main text for more information.

Table S4: Statistical comparison of all the models from BioGeoBears. LnL: log-likelihood, model parameters: d: dispersal, e: extinction, j: founder-event speciation or jump dispersal. AICc: Akaike information criterion for small sample size and AICc wt: Akaike weights.

| **Models** | **LnL** | **Number of parameters** | **d** | **e** | **j** | **AICc** | **AICc wt** |
|---|---|---|---|---|---|---|---|
| DEC | -37.18 | 2 | 0.014 | 1.00e-12 | 0 | 78.61 | 0.29 |
| **DEC+*j*** | **-35.93** | **3** | **0.01** | **1.00e-12** | **0.015** | **78.37** | **0.33** |
| DIVALIKE | -37.75 | 2 | 0.018 | 1.00e-12 | 0 | 79.75 | 0.17 |
| DIVALIKE+*j* | -36.61 | 3 | 0.012 | 1.00e-12 | 0.014 | 79.72 | 0.17 |
| BAYAREALIKE | -48.36 | 2 | 0.011 | 0.033 | 0 | 101 | 4.10e-06 |
| BAYAREALIKE+*j* | -38.09 | 3 | 0.0085 | 1.00e-07 | 0.022 | 82.68 | 0.038 |

Table S5: Divergence time and BioGeoBears Model summary on MCC tree for select nodes.

| Node | Clade | Median age (95% HPD) | DEC | DEC+*j* | DIVA | DIVALIKE+*j* | BAYAREALIKE | BAYAREALIKE+*j* |
|---|---|---|---|---|---|---|---|---|
| 53 | *Miliusa* crown | 11.85 (15.05-9.03) | B:31.37; AB:62.38 | B:40.87; AB:55.63 | B:94.96 | B:92.13 | B:36.63; AB:62.89 | B:99.72 |
| 54 | Clade C and Clade D | 10.82 (13.96-8.1) | B:92.18 | B:96.44 | B:99.96 | B:99.94 | B:82.67; AB:16.7 | B:100 |
| 55 | Clade C | 7.07 (9.91-4.75) | B:89.15; BC:10.85 | B:97.87 | B:99.95 | B:99.94 | B:98.37 | B:100 |
| 65 | Clade D | 8.05 (11.01-5.51) | B:78.73; BC:21.26 | B:92.03 | B:97.47 | B:98.17 | B:92.73 | B:99.95 |
| 66 | Clade D Indo-Burma and Wallacea, and Sahul | 5.02 (7.82-2.77) | B:99.62 | B:99.56 | B:100 | B:100 | B:97.05 | B:100 |
| 72 | Clade D Wallacea and Sahul subset | 5.38 (7.85-3.23) | BC:100 | B:68.16; BC:30.19 | BC:100 | B:44.22; BC:54.3 | B:60.94; BC:37.91 | B:97.75 |
| 73 | Wallacea and Sahul | 3.81 (5.84-2.13) | C:100 | C:100 | C:100 | C:100 | C:90.36 | C:100 |
| 77 | Clade A and B | 9.65 (12.44-7.24) | B:24.79; AB:72.21 | B:36.75; AB:59.42 | B:91.95 | B:89.61 | AB:97.21 | B:99.43 |
| 78 | Clade B | 8.51 (11.02-6.39) | A:11.79; AB:88.21 | B:28.52; A:16.98; AB:54.5 | AB:97.09 | B:58.44; A:10.3; AB:31.26 | AB:97.73 | B:95.2 |
| 79 | Clade B subset | 7.84 (10.11-5.83) | AB:94.42 | B:51.25; AB:43.11 | B:96.51 | B:93.86 | AB:96.72 | B:97.63 |
| 80 | Clade B2 | 6.65 (8.75-4.81) | B:100 | B:100 | B:100 | B:100 | B:96.08 | B:100 |
| 92 | Clade B1 | 7.02 (9.18-5.12) | AB:100 | B:45.9; AB:45.06 | AB:100 | B:68.1; AB:26.4 | AB:92.27 | B:95.48 |
| 93 | Clade B1 PI | 6.53 (8.61-4.71) | A:100 | A:100 | A:100 | A:100 | A:93.76 | A:100 |
| 102 | Clade A | 8.71 (11.82-5.82) | B:72.93; A:10.33; AB:16.74 | B:69.58; AB:21.06 | B:95.51 | B:93.76 | AB:97.66 | B:99.53 |

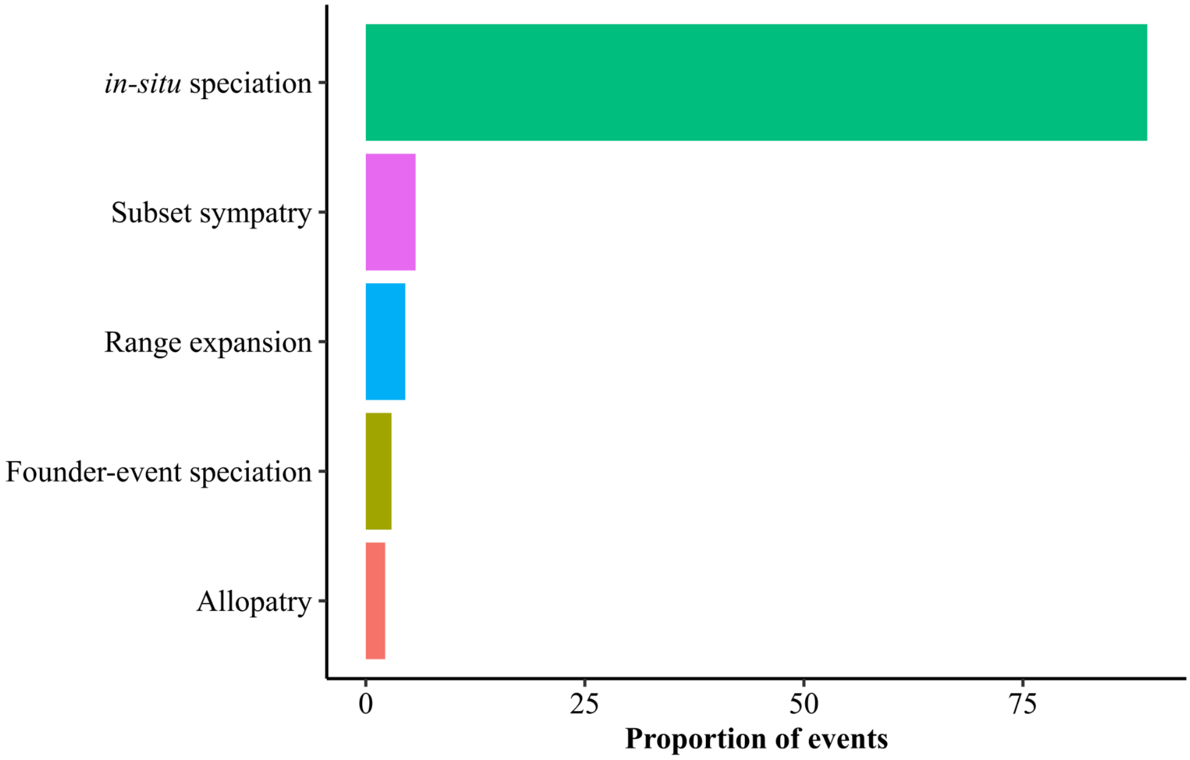


Figure S2. Proportion of overall biogeographic events as estimated from 2,000 events (100 trees sampled from the posterior and 20 BSMs per tree).

Table S6. Summary of species richness (SR), phylogenetic diversity (PD), mean pairwise distance (MPD) and their respective standardised effect sizes (ses), PDses and MPDses, speciation rates as estimated at the tips using ClaDS and DR estimated across 100 trees sampled from the posterior. Additionally, the mean biogeographic events across 2,000 events (100 trees and 20 BSMs) have been summarised.

| **Biogeographic area** | **SR** | **PD (mean ± SD)** | **MPD (mean ± SD)** | **PD.ses (mean ± SD)** | **MPD.ses (mean ± SD)** | **ClaDS (mean ± SD)** | **DR (mean ± SD)** | ***in-situ* speciation** | **Subset sympatry** | **Allopatric speciation** | **Range expansion** | **Founder-event** |
|---|---|---|---|---|---|---|---|---|---|---|---|---|
| Peninsular India (A) | 13 | 84.32 ± 11.37 | 15.05 ± 2 | -1.33 ± 0.81 | -7.44 ± 0.99 | 0.18 ± 0.04 | 0.182 ± 0.06 | 18.86 | 0.74 | 0.66 | 1.78 | 0.83 |
| Indo-Burma (B) | 35 | 169.96 ± 22.07 | 19.75 ± 2.5 | -2.56 ± 0.64 | -1.6 ± 1 | 0.21 ± 0.04 | 0.28 ± 0.16 | 64.04 | 2.13 | 1.12 | 0.50 | 0.57 |
| Wallacea and Sahul (C) | 8 | 45.25 ± 5.63 | 13.85 ± 1.78 | -3.87 ± 0.53 | -5.75 ± 1.19 | 0.21 ± 0.04 | 0.29 ± 0.12 | 8.07 | 0.03 | 0.46 | 2.34 | 1.59 |

Table S7. Number of occurrences and mean±SD of climatic values of *Miliusa* species occurring in PI.

| Sno | Species | Occurrences | Precipitation seasonality | Mean Annual Precipitation | Elevation |
|---|---|---|---|---|---|
| 1 | *Miliusa gokhalaei* | 14 | 117.85±10.71 | 3398±751.09 | 876.36±288.68 |
| 2 | *Miliusa indica* | 7 | 57.46±5.24 | 1692.14±614.81 | 768.71±438.1 |
| 3 | *Miliusa malnadensis* | 3 | 140.53±7.38 | 5245.67±889.17 | 1302.67±348.07 |
| 4 | *Miliusa cf montana* | 6 | 67.39±10.21 | 2053±105.67 | 1457±220.6 |
| 5 | *Miliusa nilagirica* | 9 | 135.9±8.08 | 3788.89±924.19 | 924.33±220.67 |
| 6 | *Miliusa sahyadrica* | 1 | 53.35±NA | 1669±NA | 333±NA |
| 7 | *Miliusa sp1* | 6 | 81.26±6.57 | 999.67±390.13 | 862.67±274.42 |
| 8 | *Miliusa tirunelvelica* | 3 | 58.04±3.93 | 2561.33±183.51 | 1338.33±18.18 |
| 9 | *Miliusa tomentosa* | 12 | 105.65±27.14 | 1292.17±723.79 | 559.75±296.77 |
| 10 | *Miliusa velutina* | 4 | 120.01±18.98 | 2173.5±957.3 | 381±320.44 |
| 11 | *Miliusa wayanadica* | 4 | 126.07±4.51 | 3613.5±492.89 | 1197.25±194.54 |
| 12 | *Miliusa wightiana* | 2 | 53.41±1.76 | 2091.5±31.82 | 1017.5±150.61 |
| 13 | *Miliusa sp2* | 3 | 87.92±7.65 | 921±257.05 | 290.67±240.19 |
